# Canary in the Forest? – Tree mortality and canopy dieback of western redcedar linked to drier and warmer summer conditions

**DOI:** 10.1101/2023.01.11.522134

**Authors:** R.A. Andrus, L.R. Peach, A.R. Cinquini, B. Mills, J.T. Yusi, C. Buhl, M. Fischer, B.A. Goodrich, J.M. Hulbert, A. Holz, A.J.H. Meddens, K.B. Moffett, A. Ramirez, H.D. Adams

## Abstract

Tree mortality and partial canopy dieback are increasing in many forest ecosystems from unfavorable climate conditions. Examining how tree growth and mortality are affected by climate variability can help identify proximate causes of tree mortality and canopy dieback. We investigated anomalously high mortality rates and partial canopy dieback of western redcedar (*Thuja plicata*, WRC), a culturally, ecologically, and economically important species in the Pacific Northwest (USA), using tree-ring methods. We sampled trees in three tree status groups—no canopy dieback, partial canopy dieback, and trees that died (0-30 years ago)—from 11 sites in coastal (maritime climate) and interior (continental climate) populations of WRC trees. In our study, WRC tree mortality was portended by on average 4-5 years of declining radial growth. Warmer and drier climate conditions in May and June that extend the annual July-September dry season reduced radial growth in 9 of 11 sites (1975-2020). Defining drought events as warm, dry May-June climate, we found that WRC trees recovered radial growth to pre-drought rates within three years when post-drought climate conditions were average or cooler and wetter than average. However, radial growth recovery from drought was slower or absent when conditions were warmer and drier during the post-drought recovery period, which appeared to lead to the widespread mortality event across coastal populations. Annually resolved tree mortality in coastal populations predominately occurred in 2017-2018 (80% of sampled trees) and coincided with exceedingly hot temperatures and the longest regionally dry period for May to September (1970-2020). In interior populations, tree mortality was associated with warmer, drier conditions from August to September. Our findings forewarn that a warming climate and more frequent and severe seasonal droughts will likely increase the vulnerability of WRC to canopy dieback and mortality and possibly other drought-sensitive trees in one of the world’s largest carbon sinks.

## INTRODUCTION

Tree mortality and canopy dieback are increasing in many forest biomes (McDowell et al. 2020, Hammond et al. 2022). The amount of canopy loss and mortality can vary widely from short-term disturbances with low-to high-severity mortality at the stand-scale (e.g., wildfire) to gradual shifts in forest demographics over decades across large forested areas (Harmon and Bell 2020, Furniss et al. 2020). More severe and frequent drought and heat events are key drivers of recent forest change (Sommerfeld et al. 2018, Anderegg et al. 2019), often interacting with other agents (e.g., bark beetle outbreaks and fire; Seidl et al. 2017). Climate change models predict continued increases in temperature and shifts in drought characteristics (e.g., more frequent, or intense) during some seasons that are expected to negatively impact forest productivity and tree survival in many but not all forest biomes (Babst et al. 2019, Hammond et al. 2022). Tree mortality is an essential and natural component of forest ecosystems, but notable losses of forest cover have substantial implications for stand structure and composition (Anderegg et al. 2012), climate regulation (Bonan 2008, Pan et al. 2011), water cycles (Mikkelson et al. 2013), and culture (Morris et al. 2018, Armstrong et al. 2022). For example, forests of the Pacific Northwest (PNW) region of western North American (including states of Washington, Oregon, and Idaho, USA and the province of British Columbia, Canada) play an outsized role in global climate regulation, because they sequester and store large quantities of carbon (Waring and Franklin 1979, Buotte et al. 2020). In forested regions where no dieback events are reported (e.g., PNW; Hammond et al. 2022), new canopy dieback events associated with tree species expected to be susceptible to drought-caused dieback may help foreshadow broader-scale forest vulnerability to climate change (i.e., canary in the coal mine).

Tree mortality can result from individual abiotic and biotic stressors or the interaction among multiple stressors (e.g., the ‘disease-decline spiral’ or ‘mortality spiral’ of Franklin et al. 1987, Manion 1991), complicating the identification of proximate causes and increasing the difficulty of prediction (Allen et al. 2015, Trugman et al. 2021). Warmer and drier climate conditions that reduce soil moisture availability, increase atmospheric drought, and elevate heat stress may push trees toward critical physiological limits for survival (McDowell et al. 2022).

Failure of hydraulic systems and dehydration of tissues may result in partial canopy dieback and mortality if unfavorable conditions persist (Adams et al. 2017, Arend et al. 2021). Elevated tree stress also increases tree susceptibility to attack by lethal biotic agents, such as bark beetles (Raffa et al. 2008), though not all tree species have known lethal biotic agents. To persist through global climate change, most trees will need to survive multiple drought or heat events resulting from the interannual variability in climate and potentially exacerbated by rising temperatures (McDowell et al. 2022, Hammond et al. 2022). Developing long-term climate change adaptation strategies requires a stronger understanding of the proximate causes of mortality and canopy dieback (abiotic versus biotic agents) at multiple spatial and temporal scales (Hennon et al. 2020).

In temperate forests, annual tree growth rings record the response to multiple environmental conditions, such as climate variability, over decades to centuries and are helpful for identifying potential causes of partial canopy dieback and mortality (Camarero et al. 2015, Hennon et al. 2020). Prolonged decreases in tree radial growth can result from slowly increasing environmental stress, such as shading or decadal shifts toward unsuitable climate conditions. In contrast, abrupt termination of growth is more commonly caused by disturbances (e.g., bark beetles or fire) or acute climate events (Cailleret et al. 2017). To forecast the implications of global climate change for tree growth, the response of radial growth to the interannual climate variability may help identify climate conditions unfavorable for tree survival (Bigler and Bugmann 2004). However, coexisting trees of the same species within a population differ in their response to stressors and susceptibility to canopy dieback or mortality (Bigler et al. 2004, Camarero et al. 2015), depending on biotic and abiotic factors (Bottero et al. 2017, Serra-Maluquer et al. 2021). For many conifer tree species, trees that died following a drought exhibited a legacy of reduced growth for years to even decades prior to death compared to surviving trees (Cailleret et al. 2017, 2019, Puchi et al. 2021) and surviving trees grew slower post-drought compared to pre-drought (i.e. low resilience; Lloret et al. 2011, DeSoto et al. 2020). Following tree death, the calendar year of tree death can be estimated with the last partial tree ring and related to the interannual variability in daily and seasonal climate conditions to determine proximate climate causes of mortality (Bigler and Rigling 2013). However, declines in growth from reduced physiological function, or subsequent mortality, can lag behind droughts (Anderegg et al. 2013, Trugman et al. 2018). Retrospective analyses of tree-rings that examine the response of forest demographic processes, such as growth and mortality, are needed to assess species’ vulnerabilities to shifting climate conditions (Lévesque et al. 2013, Clark et al. 2016), particularly for taxa and geographies of ecological, global, and cultural significance (e.g., Bradshaw et al. 2020, Case et al. 2021).

Tree species in the Cupressaceae family diversified during the rapid warming, drying climate in the Cenozoic era (rapid in geological but not anthropogenic terms) and are widely distributed across habitats ranging from coastal temperate rainforests to some of the Earth’s driest forested regions (Pittermann et al. 2012). Some of the largest, longest-lived, most resilient tree taxa on Earth are in Cupressaceae. However, members of the Cupressaceae family are being impacted by the direct (e.g., extreme drought) and indirect (e.g., fire) climate change-related stressors (e.g., Amoroso et al. 2015, Holz et al. 2020, Kannenberg et al. 2021), with potentially large implications for the variety of ecosystems they inhabit.

Within the Cupressaceae, western redcedar (*Thuja plicata* Donn ex D. Don; hereafter, WRC) has experienced anomalously high mortality and partial canopy dieback (hereafter, dieback) throughout much of its distribution within the last decade (WA DNR 2020, Goodrich et al. 2022). Recent WRC tree mortality has greatly exceeded the previously reported annual mortality rate of < 1.5% per year (Moore et al. 2003, Larson and Franklin 2010). Canopy dieback has generated concern among indigenous peoples, public land managers, private commercial foresters, and small forest landowners, because WRC trees are a culturally, economically, and ecologically critical species in the globally important forests of the PNW (Hebda and Mathewes 1984, Klinka et al. 2009, Sutherland et al. 2016). With respect to biotic agents, no known native, lethal bark beetle species are recognized as primary drivers of WRC tree mortality (USDA 2010), and WRC trees are more tolerant to many root pathogens compared to co-occurring conifer species (Morrison et al. 2014). As such, longer and hotter summer drought conditions are hypothesized as the likely cause of the recent WRC tree dieback in the PNW (Seebacher 2007, Kral et al. 2020), with specific climate effects likely varying between interior populations with drier, continental climates and coastal populations with more mesic, maritime climate. Compared to its common tree associates in the PNW region (e.g., Douglas-fir, *Pseudotsuga menziesii* Franco), WRC is an understudied species and no peer-reviewed studies to date have examined WRC tree canopy dieback.

We investigated how WRC tree radial growth and dieback responded to climate variability and drought in coastal and interior populations in the PNW (11 sites). To identify characteristics of radial growth that may relate to higher likelihood for survival, co-existing WRC trees were sampled in three health-status groups based on canopy condition in summer 2021: no canopy dieback (<40% canopy dieback; hereafter, *healthy*), partial canopy dieback (>40% and <100% canopy dieback; hereafter, *unhealthy*), and trees that were found dead and presumably died in the last three decades (no green canopy; hereafter, *dead*). We addressed four research questions. (1) How did the radial growth rates of unhealthy or dead trees differ from those of neighboring healthy trees? We expected that unhealthy and dead trees would experience several years of reduced tree growth prior to death or 2020 (the last full ring sampled from unhealthy and healthy trees) compared to healthy trees. (2) How did radial growth respond to the interannual variability in climate conditions from 1975 to 2020 (or mortality date) and how did these climate-growth relationship vary by tree status and population? We expected that reduced radial growth would be correlated with below-average soil moisture availability as indicated by low spring and summer precipitation and high spring and summer temperature (Brubaker 1980) and that unhealthy and dead trees would be more strongly related to climate variability than healthy trees (Cailleret et al. 2019). (3) When pre-drought conditions and drought severity were similar, did radial growth responses to drought differ between warm/dry and cool/wet post-drought conditions during the recovery period, and did this relationship vary by tree status and population? We expected that warmer, drier climate conditions in the post-drought period (three years) would increase the time required to reach pre-drought growth rates and reduce post-drought compared to pre-drought growth rates (i.e., resilience; Lloret et al. 2011), especially for unhealthy and dead trees (i.e., differential post-drought resilience). No difference was expected in the reduction in radial growth during the drought year compared to pre-drought (i.e., resistance; Lloret et al. 2011), because drought severity was similar. (4) How was annually-resolved tree mortality affected by interannual variability in weather and climate conditions? We expected that tree mortality would be associated with below-average soil moisture availability as indicated by anomalously warm and dry spring and summer climate conditions. Our results provide key insights into the weather and climate effects on recent WRC tree dieback and the implications of climate change for WRC tree populations.

## METHODS

### Study Area

We studied WRC tree dieback in the coastal and interior populations in the northwestern US (Table 1, S1; Fig. 1). Similar to other Cupressaceae conifers, WRC trees inhabit a broad gradient in annual precipitation from temperate rainforests (> 500 cm yr^-1^) of the coast and windward slopes of the Cascade and Olympic Mountains to considerably drier rainshadows in coastal forests and dry interior forests (∼70 cm yr^-1^) (Minore 1983). In interior populations (elevation range, 800-2130 m), WRC trees grow best on riparian or wet sites at lower elevations and all mid-elevation hillslopes, whereas in coastal populations WRC trees are more widely distributed across their elevation range (sea-level to ∼2000 m; Minore 1983). Both populations experience a relatively wetter and cooler period (October to June) followed by an extended warm and dry summer season (July to September) during which soil moistures drop precipitously based on soil characteristics (Franklin and Dyrness 1973, Baker et al. 2019). Relative to the more moderate, maritime climate characteristic of the coastal WRC population, the continental climate of the interior WRC population typically spans a wider annual range of conditions, with notably colder and snowier winters, warmer and drier summers, and greater summer moisture deficits (Michalet et al. 2021). During the 1895-2020 period, mean annual air temperatures in the Pacific Northwest warmed by ∼1.1°C, with a majority of warming occurring from mid-1980s to 2020, and there was no long-term trend in annual precipitation (Kunkel et al. 2022). Average annual air temperatures are expected to warm 1.9°C (RCP 2.6) to 3.1°C (RCP 8.5) by the 2050s in the PNW compared to 1950-1999 (Rogers and Mauger 2021). Annual precipitation is expected to remain stable, but summer precipitation is expected to decrease by the mid and late 21^st^ century (Dalton and Fleishman 2021).

**Table 1:** Characteristics of the 11 sampled sites, including elevation (Elev.), stand basal area (BA), estimated stand initiation (< 150 or > 150 years; based on tree ages in Fig. S1), modeled site climate moisture deficit (CMD; 1980-2010 normals Wang et al. 2012), and number of sampled and successfully crossdated trees by tree health status group (healthy, unhealthy, dead). Stand basal area was estimated by averaging estimates of basal area at each sampled tree.

| Pop. | Site | Elev<br>(m) | Stand<br>BA<br>(m<sup>2</sup>/ha) | Stand<br>init.<br>(years) | CMD<br>(mm) | No. trees sampled<br>(crossdated) |  |  |
| --- | --- | --- | --- | --- | --- | --- | --- | --- |
|  |  |  |  |  |  | Healthy | Unhealthy | Dead |
| Interior | RG | 1080 | 34.8 | >150 | 160 | 10 (10) | 10 (10) | 0 |
|  | HG | 1315 | 51.0 | >150 | 250 | 11 (11) | 11 (11) | 0 |
|  | SL | 715 | 28.8 | <150 | 350 | 4 (4) | 6 (6) | 5 (5) |
|  | MP | 1034 | 29.4 | <150 | 420 | 10 (10) | 10 (10) | 0 |
|  | LI | 762 | 37.1 | <150 | 500 | 10 (9) | 10 (9) | 9 (9) |
| Coastal | B | 308 | 25.6 | <150 | 195 | 10 (10) | 10 (10) | 10 (10) |
|  | L | 158 | 26.6 | <150 | 240 | 10 (10) | 10 (9) | 10 (8) |
|  | IC | 875 | 32.4 | >150 | 300 | 10 (10) | 10 (9) | 0 |
|  | TR | 140 | 25.3 | <150 | 340 | 10 (10) | 10 (10) | 10 (10) |
|  | RF | 59 | 17.7 | <150 | 420 | 10 (10) | 10 (10) | 10 (10) |
|  | M | 202 | 23.0 | <150 | 430 | 10 (10) | 10 (10) | 10 (8) |

Site names abbreviations: Roosevelt Grove (RG), Hobo Grove (HG), Slate (SL), McCroskey Park (MP), Lindsey (LI), Barlow Wayside (BW), Little Mountain (LM), Icicle Creek (IC), Trillium (TR), Ramirez Farm (RF), Mount Richmond (MR)

**Fig. 1:**
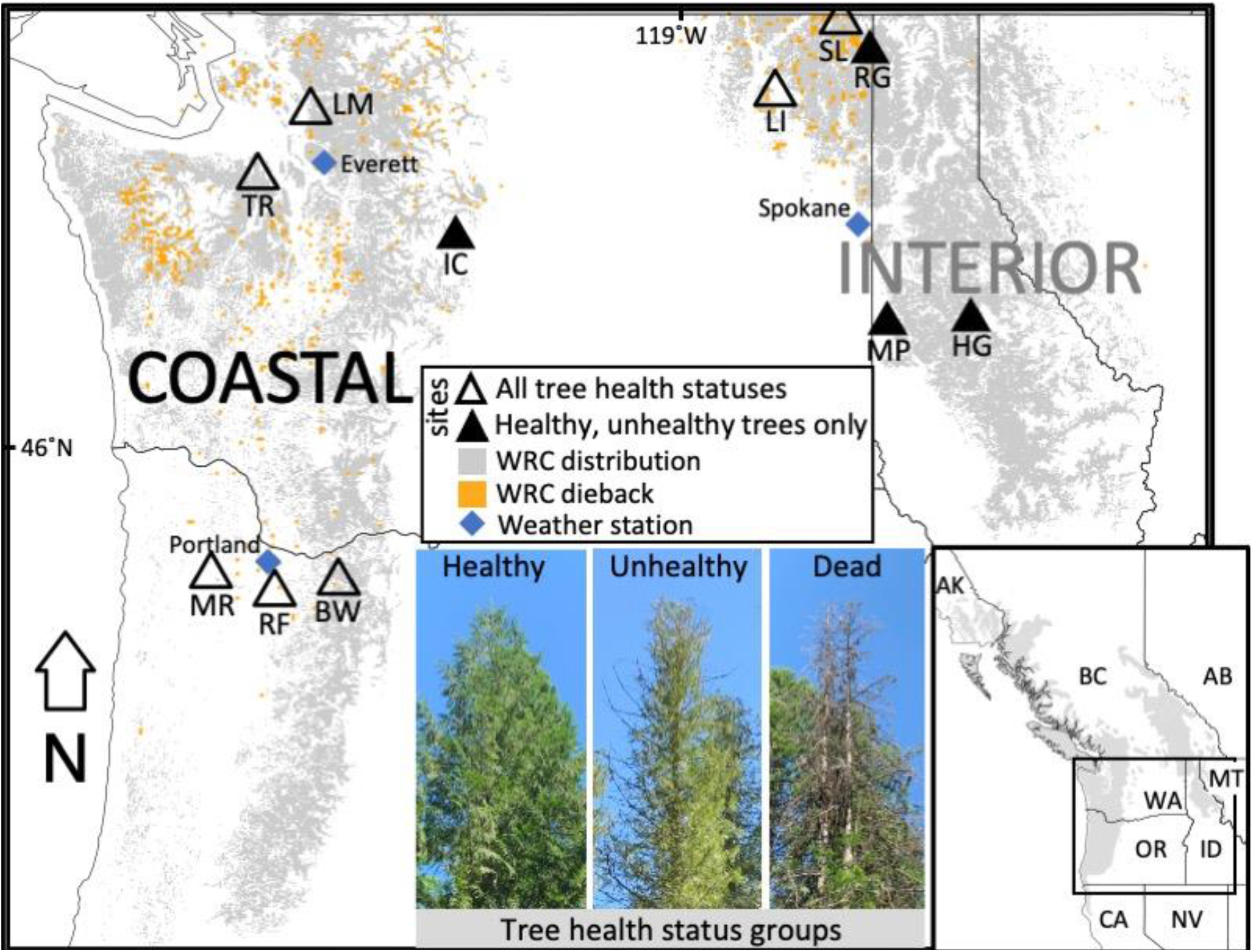
Locations of the 11 sampled sites (triangles) with western redcedar dieback and mortality (orange; USDA Forest Service 2021) within the statistically modeled distribution of western redcedar (grey; *Thuja Plicata;* Ellenwood et al. 2015) in the northwestern United States (map) and western North America (map inset; (Little 1971)). Photos show examples of trees in three tree health status categories: healthy (<40% canopy dieback), unhealthy (> 40% and < 100% canopy dieback), and dead (no green canopy).

Traits that contribute to WRC’s broad distribution and long lifespan (> 1000 years) include high rates of survival under unfavorable conditions (e.g., high shade, nutrient-poor soils, and wet conditions), high resistance to biotic agents of mortality (due to high concentration of terpenes), and slow growth (i.e., dense wood) (Antos et al. 2016). WRC trees occupy a sub-dominant canopy position in many mixed-conifer communities today, but it can dominate and form pure stands at some sites, especially in wet or nutrient-poor soils (Kranabetter et al. 2013). Common tree-species associates of WRC trees vary from wetter forests of western hemlock (*Tsuga heterophylla* (Raf.) Sarg.), Sitka spruce (*Picea sitchensis* (Bong.) Carrière), and red alder (*Alnus rubra*) to drier forests of Douglas-fir, grand fir (*Abies grandis* (Dougl. ex D. Don) Lindl.), and bigleaf maple (*Acer macrophyllum* Pursh) (Minore 1983). Since European colonization, nearly all forests within WRC’s distribution have been heavily managed by public agencies and private timber companies for timber extraction and some portions have been converted to agricultural and urban areas.

### Field Sampling

In summer 2021, we sampled WRC trees in 11 sites, six in coastal populations and five in interior populations (Fig. 1; Table 1). To identify suitable areas for sampling, we relocated in the field areas previously mapped as WRC dieback by Goodrich et al. (2002) and selected sampling areas to span a range in climatic moisture deficit (CMD) and climate conditions (Table 1, S2).

CMD is the annual total of monthly atmospheric evaporative demand not met by monthly precipitation and is a proxy for both atmospheric and soil moisture limitations (Wang et al. 2012). Potential sites were excluded if we observed evidence of recent forest management (e.g., thinning in the last 30 years), recent disturbance (e.g., wind, fire), or human development.

In an area with WRC dieback, we used a targeted sampling approach to identify 10 neighboring triplets of healthy, unhealthy, and dead trees or pairs when dead trees were not present at the site (20-30 trees per site) (Table 1, Fig. 1). Tree health-status groups were defined as the percent of skylight visible through the live, normally fully-foliated portion of the crown (i.e., estimate of dieback), following similar recent studies (see Fig. 1 for examples) (Camarero et al. 2015, Comeau et al. 2019). At each site, triplets (or pairs) were matched for similar tree size (DBH, diameter at breast height) and tree height, and all trees were within a qualitatively similar abiotic (e.g., aspect and site moisture availability) and biotic (e.g., stand basal area) environment.

For each tree, we extracted two tree cores (increment core diameter: 0.5mm) at breast height (∼1.4 m) with the bark and outer most ring intact. Then, we recorded the following tree attributes: DBH, height, status (healthy, unhealthy, dead), canopy position (dominant, co-dominant, intermediate, suppressed), percent canopy dieback (0-100% in 10% classes), tree crown symptoms (top kill, branch dieback, thinning crown), and pest and pathogen damage (e.g., bark beetles, root disease, heartwood decay). We observed no evidence of insect damage in the cambium of dead WRC trees. Additionally, no consistent biotic damage agent of mortality was found in 148 field site visits with WRC tree dieback in Washington and Oregon (Goodrich et al. 2022), though no roots were destructively sampled for pathogens. The sampled area differed by site (∼0.5-2 ha) due to the density and extent of mortality and partial canopy dieback symptoms. Topographic descriptors, location, and structure and composition were recorded at each site (Table S1).

### Tree core processing and constructing tree-ring chronologies

We processed WRC tree cores following standard dendroecological procedures (Speer 2010). Cores were mounted, sanded, aged, visually crossdated, and scanned (2400 dpi with Epson Expression 12000XL). Scans of cores were measured with a precision of 0.01 mm (CooRecorder; Maxwell and Larsson 2021). For each site, we quantitatively crossdated both cores from 10 healthy and 10 unhealthy trees using marker rings and time-shifted correlation coefficients (‘dplR’ package in R; Bunn 2010, R Core Team 2016). Then, ring-width series from dead trees were crossdated by comparing to crossdated ring-width series from healthy and unhealthy trees at the same site. The year of tree death was estimated as the year of the outermost partial or full ring (Bigler and Rigling 2013). If the year of the outermost rings differed for a pair of cores from a dead tree (e.g., due to partial cambial dieback prior to death), we assumed that the tree died in the year of the most recent full or partial ring. Cores with missing outermost rings (weathered) and cores that could not be confidently crossdated were excluded from further analysis. We successfully crossdated 90% of all the sampled cores (493 of 547 cores), together representing 94% of the sampled trees (263 of 281; Table 1). To achieve per-tree radial growth data for analysis, the two ring-width measurements from an individual tree were averaged for any overlapping years and non-overlapping years were excluded (Adams and Kolb 2005). We calculated basal area increment (BAI) using raw ring-width measurements and the tree’s DBH (Biondi 1999).

To test climate-growth relationships, we constructed tree ring-width chronologies for each site from 1975 to 2020, a period with > 15 trees per site in the chronology (Fig. S1). Ring-width series were detrended using a cubic smoothing spline (50% frequency cutoff, 30-year wavelength; Klesse 2021). Spline wavelengths +/- 10 years in length produced similar results. Detrending was necessary to account for the decline in absolute ring-width associated with increasing tree bole diameter and retain intra-series annual to decadal variability (e.g., the higher-frequency ring-width signal). Temporal autocorrelation in detrended ring-width series was removed with autoregressive models (i.e., prewhitening function in ‘dplR’ package). Ring-width series were averaged using the Tukey’s biweight robust mean (unaffected by outliers) to construct residual site chronologies for all trees and each tree status group (Fig. S2; ‘chron’ function in ‘dplR’ package). For the site tree-ring chronologies, we report mean inter-series correlation (i.e., MSI, the correlation among all ring-width series) and expressed population signal (i.e., EPS, variance explained by the sample trees) of detrended ring-width series (Wigley et al. 1984).

### Climate and Drought Datasets

To characterize drought conditions, we used the standardized precipitation evaporation index (SPEI), a multi-scalar drought index indicative of soil moisture availability (Vicente-Serrano et al. 2010). We extracted monthly precipitation and air temperature data for each year from the 4-km grid cell containing plot center (PRISM 2022) and calculated SPEI by month, season, and water year (October to September) from 1900 to 2020 (‘spei’ package in R; Begueria and Vicente-Serrano 2017). To assess the possible influence of weather conditions on tree mortality, we downloaded instrumental daily weather data for three representative locations within the study area (Fig. 1) in northwestern Washington State (Everett, WA; station USW00024222), northeastern Washington State (Spokane, WA; station USW00024157), and northwestern Oregon state (Portland, OR; station USW00024229; NOAA 2022). For each year from 1970 to 2020, we computed the following parameters from May to September: (1) maximum temperature index (MTI; the number of days that monthly maximum daily air temperature exceeded the 90th percentile) and (2) the dry period index (DPI; the longest consecutive period of days with < 1.0 mm of precipitation).

### Statistical Analysis

#### Tree radial growth preceding tree death and during canopy dieback (Question 1)

To understand how radial growth rates of unhealthy and dead trees differed from healthy trees, we calculated the annual growth ratio (30 most recent years) for co-located pairs of 1) dead and healthy and 2) healthy and unhealthy trees in each site (Cailleret et al. 2017). Ratios were calculated annually per tree pair by dividing a healthy tree’s BAI by a dead or unhealthy tree’s BAI. Annual growth ratios less than one indicate that healthy trees grew more in that year than dead or unhealthy trees. Although an effort was made in the field sampling to control for size effects on tree growth, the matching was not perfect, and to maximize available samples for each pair, the DBH of the healthy tree was within 7 cm DBH of the unhealthy or dead tree (41 tree pairs for dead-healthy, 56 tree pairs for unhealthy-healthy). As annual growth ratios were similar among sites within populations, we computed and present population averages and 95% confidence intervals of annual growth ratios for interior and coastal populations (‘boot’ package in R; Angelo and Ripley 2021). Heartwood decay and core collection height precluded estimation and incorporation of tree age. We assumed all mortality was the consequence of the same process as no available evidence indicated otherwise (e.g., bark beetle galleries, root disease); however, we estimated that dead WRC trees were all < 150 years old and thus a fraction of their potential age (which can exceed 1500 years; Daniels 2003), suggesting a minimal effect of age on our results.

#### Tree radial growth response to interannual climate variability (Question 2)

Tree radial growth responses to climate variability can help identify past climate conditions that increased tree stress and may predispose trees to mortality. Because warming and overall atmospheric aridity trends increased significantly across the western US, including the PNW, starting in the year 2000 (e.g., Abatzoglou and Williams, 2016), we conducted separate correlation function analyses for two periods, 1975 to 1999 and 2000 to 2020. The effects of period of analysis and tree health status had negligible effects on climate-growth relationships for either period (Fig. S3-S7) and were excluded from further analysis. We tested correlations from 1975 to 2020 between tree-ring site chronologies (all tree health status groups; ‘treeclim’ package in R; (Zang and Biondi 2015)), using ring-width indices and three site climate variables: (1) monthly maximum air temperature, (2) monthly total precipitation, and (3) monthly SPEI. Correlations were tested from April of year prior to ring formation (lagged effects) to September of ring formation year for each site. We considered correlations coefficients significant at α = 0.05, and only interpret results when statistically significant results were found in two or more sites within each population (interior or coastal).

#### Tree radial growth response to drought events (Question 3)

Declines in radial growth during and after drought conditions may indicate reduced tree vigor and greater susceptibility to mortality during subsequent drought events (DeSoto et al. 2020). For each site, we compared radial growth responses to two exemplary drought years from 1975 to 2015 (period with > 5 samples in each tree health status group per site, Fig. S1). We defined drought years as May-June SPEI < -1.0 based on the important influence of May-June SPEI on annual tree radial growth on both populations (see results for Question 2). Both drought years had similar pre-drought and drought year May-June SPEI, but post-drought conditions were warm/dry or cool/wet (Fig. S8-S9). For all sites in coastal and interior populations, we selected the 2015 regional drought year (lowest or second lowest May-June SPEI from 1975 to 2020), which was followed by three years of warm/dry conditions in spring and summer (Fig. S8-S9). For the second drought year, we selected site-specific drought years, with the lowest May-June SPEI (other than 2015) at each site from 1975-2014. All second exemplary drought years occurred prior to 2004 (hereafter, pre-2004 drought year) and were followed by three years of notably cooler, wetter conditions in spring and summer (see Fig. S8 for drought years by site).

To characterize radial growth responses to drought events, we extracted three growth metrics from each tree’s detrended ring-width index to calculate four resilience indices (Lloret et al. 2011, Thurm et al. 2016, Schwarz et al. 2020). For each tree, the growth metrics were as follows: *Dr* was the mean annual growth during the drought year, *PreDr* was the mean annual growth for the three years before the drought year, and *PostDr* was the mean annual growth for the three years after the drought year. For trees that died less than three years after the drought year, we calculated PostDr from the years prior to death, excluding partial rings. We also recorded the number of years post-drought required to return to *PreDr*. For each tree and drought event, we computed the following resilience indices:

1. Resistance (Rt) = *Dr/PreDr*, (i.e., growth during drought year relative to pre-drought)
2. Recovery (Rc) = *PostDr/Dr*, (i.e., growth post-drought relative to during drought year)
3. Recovery period (Rp) = Number of years post-drought required to reach pre-drought growth rates
4. Resilience (Rs) = *PreDr/PostDr* (i.e., growth pre-drought relative to post-drought)

We selected a reference period of three years for the resistance, recovery, and resilience indices based on the average length of the recovery period (three years) following the pre-2004 drought year. Unhealthy and healthy trees (57% or 107 of 189 trees) that were still in their recovery period from the 2015 drought year in 2020 (last full ring) were assigned a recovery period of six years (i.e., the minimum period for recovery).

To test for differences in radial growth responses to the two exemplary drought years by tree status and population, we constructed one linear mixed model (LMM; Pinheiro et al. 2022) for each of four resilience indices (response variables). We tested the following predictor variables: drought event year (categorical: 2015 with warm, dry post-drought conditions or pre-2004 with cool, wet post-drought conditions), tree health status (categorical: healthy, unhealthy, dead), WRC tree population (categorical: interior or coastal), the interaction between drought event year and tree health status, and the interaction between drought event year and population. Several other predictor variables, including DBH or tree height, stand basal area (measured in the field with a basal area prism at each sampled tree to estimate competition for resources), site elevation, and site CMD, were also tested in bivariate preliminary models, but were not statistically significant predictor variables (*P* > 0.05) and were excluded from further analysis.

We included a random effect of trees within site to account for repeated measures of individual trees in two drought years and potential differences among sites in resilience indices. Model residuals were checked, and model fit was assessed using the conditional and marginal r^2^ (‘MuMin’ package in R (Bartoń 2018)).

#### Effect of climate and weather variability on tree mortality (Question 4)

To assess whether WRC tree mortality corresponded with warm/dry spring and summer climate and weather conditions, we graphically compared annually resolved WRC tree mortality dates to climate (SPEI) and weather parameters (DPI and MTI) from 1970 to 2020. For SPEI, we selected May to September SPEI for coastal populations and August to September SPEI for interior populations after comparing monthly and water year SPEI (1970-2020) to mortality years (Fig. S10).

## RESULTS

### Tree radial growth preceding tree death and during canopy dieback (Question 1)

Trees that died (dead) and partial canopy dieback trees (unhealthy) experienced a period of reduced radial growth prior to death or 2020, respectively, compared to healthy trees (Fig. 2; Fig. S11-S12). Average annual growth ratios (based on BAI) of dead trees (compared to healthy) were notably lower for four years (interior) and five years (coastal) prior to tree death (95% confidence interval < 1.0), with similar annual growth ratios for the prior ∼25 years (−30 to -5 years; Fig. 2A). However, the timing and duration of the reduction in annual growth ratios of dead compared to healthy trees was highly variable among individuals but consistent within populations (Fig. S12). Radial growth of dead trees was lowest in the last full year of growth, with an average reduction in annual growth ratios of 112% in interior (range in individuals, 0-262%) and 78% in coastal populations compared to healthy trees (range of individuals, 96-191%; Fig. 2, Fig S12).

**Fig 2:**
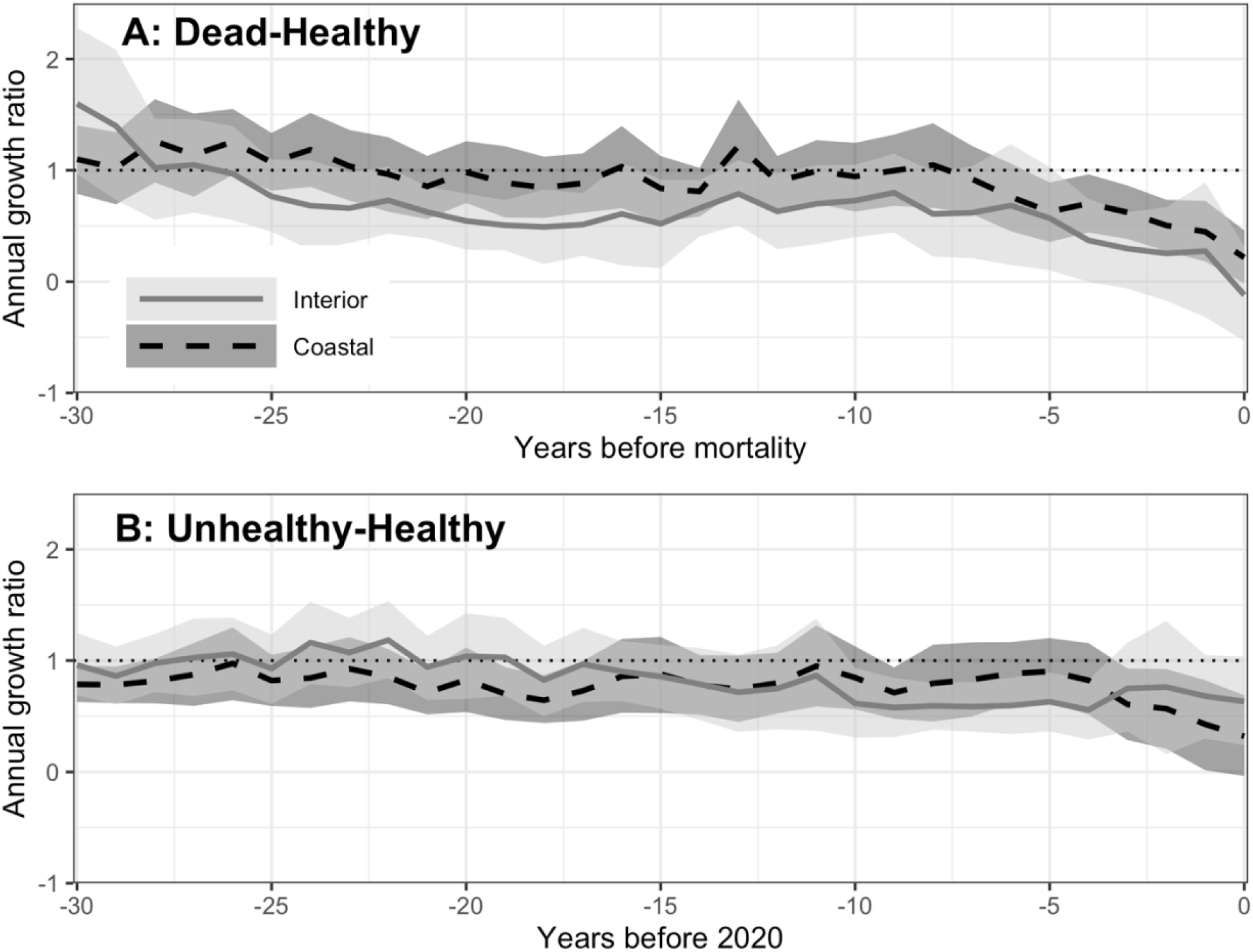
Annual growth (Basal Area Increment; BAI) ratios for **A)** dead vs. healthy tree pairs (7 sites; 41 tree pairs) and **B)** unhealthy vs. healthy tree pairs (9 sites; 56 tree pairs) computed from the most recent 30 years of BAI tree growth. The gray and dashed black lines are the means, and the light and dark gray shaded ribbons are the 95% confidence intervals of all trees in the interior and coastal populations, respectively. An annual growth ratio significantly less than one indicates that healthy trees grew more than their paired, co-located dead or unhealthy trees in that year (or the opposite, if ratio >1).

For unhealthy and healthy tree pairs, average annual growth ratios were similar for most of the 1990-2020 study period (95% confidence interval included 1.0), except 2017-2020 in coastal populations and 2009-2017 in interior populations (Fig. 2B). In coastal populations, annual growth ratios of unhealthy trees steadily declined from 2017 to 2020, with the lowest annual growth ratio in 2020 (68%; range in individuals, 90-248%). In interior populations, unhealthy trees in some sites experienced ∼40% less growth than healthy trees for multiple years in the last decade (e.g., SL, RG), while annual growth ratios in other sites were comparable (ratio near 1.0) during the study period (e.g., MP, HG; Fig. 2B, Fig. S12)

### Tree radial growth response to interannual variability in climate (Question 2)

All pairs of site tree-ring chronologies were relatively well correlated from 1975 to 2020 for sites in coastal populations on the westside of the Cascade Mountains (Spearman’s rho >0.36, *p <*0.01 for ‘all trees’, all coastal sites except IC) and sites in interior populations (Spearman’s rho >0.30, *p <*0.05 for ‘all trees’; Fig. S2, S13). The IC site (coastal population on eastside of Cascades Mountains) was more strongly correlated with sites in the interior (Spearman’s rho: 0.44-0.66) than coastal population (Spearman’s rho: 0.29-0.39).

Radial growth (ring-width index from tree-ring site chronologies) was correlated to the interannual variability in monthly maximum temperature, total precipitation, and the SPEI drought index for some months from April of the year prior to ring formation to September of the growth year (1975-2020; Fig. 3). In coastal populations, radial growth in two or more sites increased under the following conditions: cooler maximum June temperatures in growth year (negative correlation, *p* < 0.05 for 3 of 6 sites), higher total May or June precipitation in growth year (positive correlation, *p* < 0.05 for 6 of 6 sites), higher Aug SPEI in year prior to ring formation (*p* < 0.05 for 2 of 6 sites), and higher May or June SPEI in growth year (*p* < 0.05 for 6 of 6 sites; Fig. 3). In interior populations, radial growth at three interior sites (SLA, LI, MP) increased under the following conditions: warmer January (growth year) maximum temperatures (*p* < 0.05 for 3 of 3 sites), higher June SPEI in growth year (*p* < 0.05 for 2 of 3 sites), higher July SPEI in year prior to ring formation (*p* < 0.05 for 2 of 3 sites), and lower SPEI in November prior to ring formation (*p* < 0.05 in 2 of 3 sites). The two high-elevation sites with lower climate moisture deficits in interior populations were relatively unaffected by interannual climate variability (1 of 54 and 2 of 54 correlation tests with *p* < 0.05 at HG and RG, respectively).

**Fig. 3:**
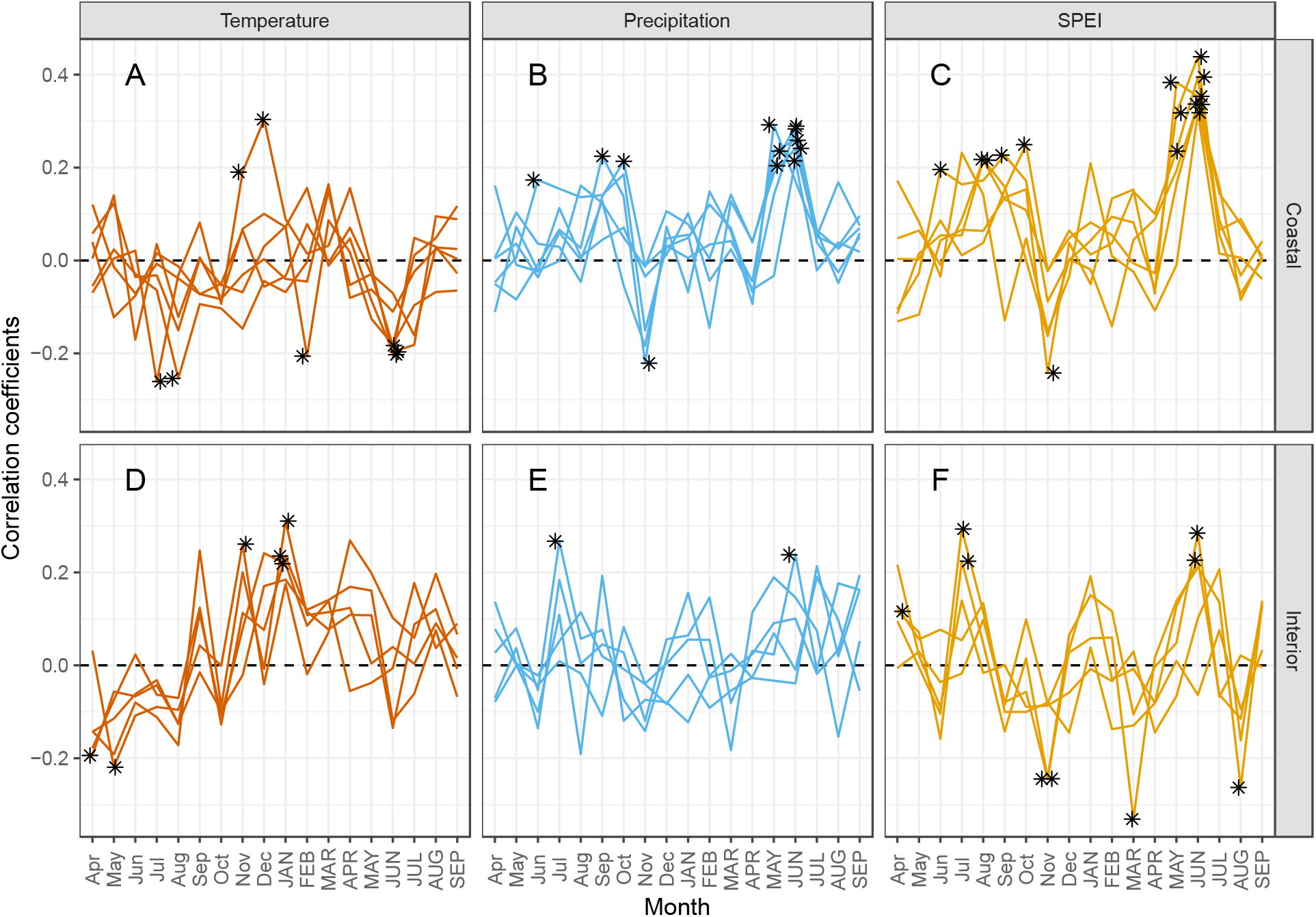
Correlation coefficients between the site tree ring chronologies and monthly climate variables, separated by coastal and interior populations of western redcedar from 1975-2020. Climate variables included monthly (A, D) maximum temperature, (B, E) total precipitation, and (C, F) Standardized Precipitation Evaporation Index (SPEI) from the year prior to growth (lower case) to the growth year (upper case). Positive values of SPEI indicate cooler, wetter conditions and negative values indicate warmer, drier conditions. Stars indicate significant relationships (*P* < 0.05). See Fig S3-S5 for minimal effect of tree status on climate-growth relationships.

### Tree radial growth response to drought events (Question 3)

Compared to one year pre- and post-drought, radial growth was lowest in the drought year for ∼50% of WRC trees in the pre-2004 (all from 1975-2004) and 2015 drought years. For trees in all health status groups, annual growth rates were 12.6% (mean, SE 1.1%) lower during the drought years compared to the average of the prior three years (i.e., resistance index, Rt; Fig. 4A-B). The resistance index (Rt) LMM did not indicate differences in resistance by drought year, tree status, population, or the interaction of drought year and tree status (Table 3). In contrast, WRC trees in the interior population were notably more resistant to the 2015 drought (mean growth reduction -8.3%, SE 1.9%) than trees in the coastal population (mean -20.2%, SE 1.5%; Fig. 4A), as demonstrated by the Rt LMM (‘Population (interior)*Drought yr. (2015)’ variable in Rt model, Table 3). However, the 2015 drought (May-June SPEI) was also slightly less severe at sites in the interior population (median -1.37) compared to the coastal population (median -1.82; Fig, S9).

**Table 2:**
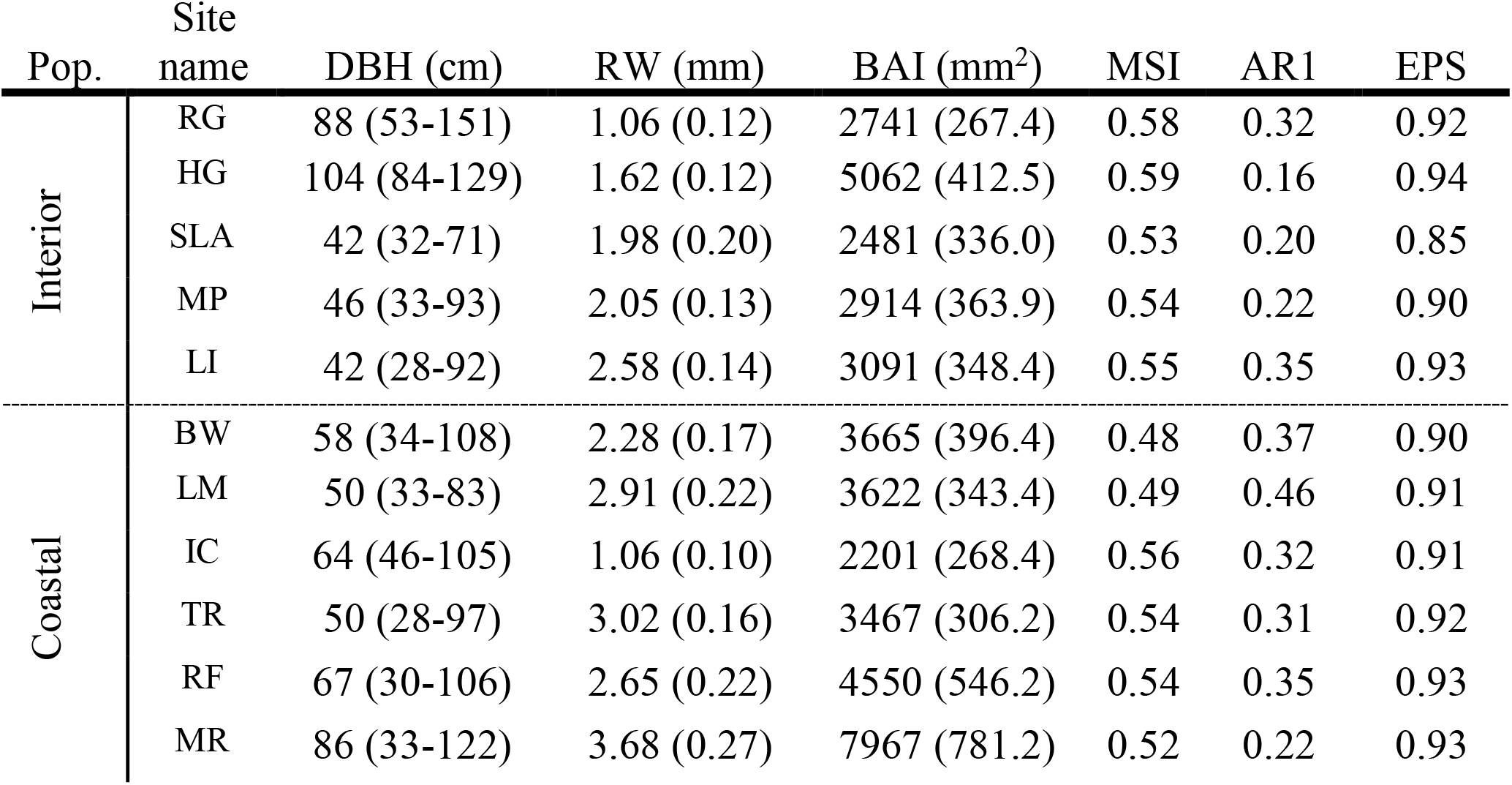
Characteristics of trees and ring-width chronologies (1975-2020) for the 11 sites: tree diameter at breast height (DBH mean and (range)), ring width (RW mean (standard error)), basal area increment (BAI mean (standard error)), mean series intercorrelation (MSI), series mean first-order autocorrelation (AR1), and expressed population signal (EPS). Each site’s analysis included data from at least 10 trees (sample depth) at the start of the chronologies in 1975 (Fig. S1).

**Table 3:** Summary of four linear mixed effects models for resistance index (Rt), recovery period (Rp), recovery index (Rc), and resilience index (Rs) of radial growth across all trees. For fixed effects, values are estimates of coefficients (β), standard error (SE), degrees of freedom (DF), and p-values (*p*). For random effects, values are standard deviations (SD) of intercept and residual error. The following predictors were tested: tree health status (healthy, unhealthy, dead), drought year (2015 or a drought prior from 1975 to 2004), WRC tree population (interior or coastal), the interaction between tree status and drought event, and the interaction between drought event and population. *p* < 0.05 were considered statistically significant.

| Response | Predictor | $\beta$ | SE | DF | t | p | SD (Residual) |
| --- | --- | --- | --- | --- | --- | --- | --- |
| **Rt** |  |  |  |  |  |  |  |
| Fixed | Intercept | 0.83 | 0.03 | 229 | 27.03 | **<0.01** |  |
|  | Drought (2015) | -0.02 | 0.02 | 201 | -0.86 | 0.39 |  |
|  | Status (dead) | -0.04 | 0.03 | 229 | -1.31 | 0.19 |  |
|  | Status (unhealthy) | 0.03 | 0.03 | 229 | 1.19 | 0.24 |  |
|  | Population (interior) | 0.06 | 0.04 | 9 | 1.44 | 0.18 |  |
|  | Drought (2015)*Status (dead) | 0.03 | 0.04 | 201 | 0.65 | 0.52 |  |
|  | Drought (2015)*Status (unhealthy) | 0.00 | 0.04 | 201 | 0.04 | 0.97 |  |
|  | Population (interior)*Drought (2015) | 0.10 | 0.03 | 200 | 2.93 | **<0.01** |  |
| Random | Site |  |  |  |  |  | 0.06 |
|  | Tree/site |  |  |  |  |  | 0 (0.17) |
| Trees/sites | 242/11 |  |  |  |  |  |  |
| R<sup>2</sup>m/R<sup>2</sup>c | 0.05/0.15 |  |  |  |  |  |  |
| **Rp** |  |  |  |  |  |  |  |
| Fixed | Intercept | 2.85 | 0.35 | 179 | 8.07 | **<0.01** |  |
|  | Drought (2015) | 1.23 | 0.35 | 179 | 3.55 | **<0.01** |  |
|  | Status (unhealthy) | -0.21 | 0.35 | 177 | -0.60 | 0.55 |  |
|  | Population (interior) | -0.20 | 0.41 | 9 | -0.47 | 0.65 |  |
|  | Drought (2015)*Status (unhealthy) | -0.16 | 0.49 | 179 | -0.32 | 0.75 |  |
|  | Population (interior)* Drought (2015) | -0.85 | 0.49 | 178 | -1.74 | 0.08 |  |
| Random | Site |  |  |  |  |  | 0.56 |
|  | Tree/site |  |  |  |  |  | 0 (0.23) |
| Trees/sites | 189/11 |  |  |  |  |  |  |
| R<sup>2</sup>m/R<sup>2</sup>c | 0.06/0.11 |  |  |  |  |  |  |
| **Rc** |  |  |  |  |  |  |  |
| Fixed | Intercept | 1.16 | 0.03 | 228 | 38.02 | **<0.01** |  |
|  | Drought (2015) | -0.15 | 0.03 | 183 | -4.75 | **<0.01** |  |
|  | Status (dead) | 0.09 | 0.04 | 228 | 2.44 | **0.02** |  |
|  | Status (unhealthy) | 0 | 0.03 | 228 | 0.03 | 0.98 |  |
|  | Population (interior) | 0.02 | 0.03 | 9 | 0.46 | 0.66 |  |
|  | Drought (2015)*Status (dead) | -0.15 | 0.05 | 183 | -2.72 | **0.01** |  |
|  | Drought (2015)*Status (unhealthy) | -0.05 | 0.04 | 183 | -1.13 | 0.26 |  |
|  | Population (interior)* Drought (2015) | -0.02 | 0.04 | 182 | -0.47 | 0.64 |  |
| Random | Site |  |  |  |  |  | 0.04 |
|  | Tree/site |  |  |  |  |  | 0 (0.20) |
| Trees/sites | 242/11 |  |  |  |  |  |  |
| R<sup>2</sup>m/R<sup>2</sup>c | 0.20/0.23 |  |  |  |  |  |  |
| **Rs** |  |  |  |  |  |  |  |
| Fixed | Intercept | 0.94 | 0.02 | 228 | 39.82 | **<0.01** |  |
|  | Drought (2015) | -0.14 | 0.03 | 178 | -5.3 | **<0.01** |  |
|  | Status (dead) | 0.06 | 0.03 | 228 | 1.67 | 0.1 |  |
|  | Status (unhealthy) | 0.03 | 0.03 | 228 | 1.25 | 0.21 |  |
|  | Population (interior) | 0.07 | 0.02 | 9 | 3.06 | **0.01** |  |
|  | Drought (2015)*Status (dead) | -0.05 | 0.05 | 178 | -1.1 | 0.27 |  |
|  | Drought (2015)*Status (unhealthy) | 0.00 | 0.04 | 178 | -0.11 | 0.91 |  |
|  | Population (interior)* Drought (2015) | 0.11 | 0.04 | 177 | 3.13 | **<0.01** |  |
| Random | Site |  |  |  |  |  | 0.02 |
|  | Tree/site |  |  |  |  |  | 0.04<br>(0.17) |
| Trees/sites | 241/11 |  |  |  |  |  |  |
| R<sup>2</sup>m/R<sup>2</sup>c | 0.19/0.23 |  |  |  |  |  |  |

**Fig. 4:**
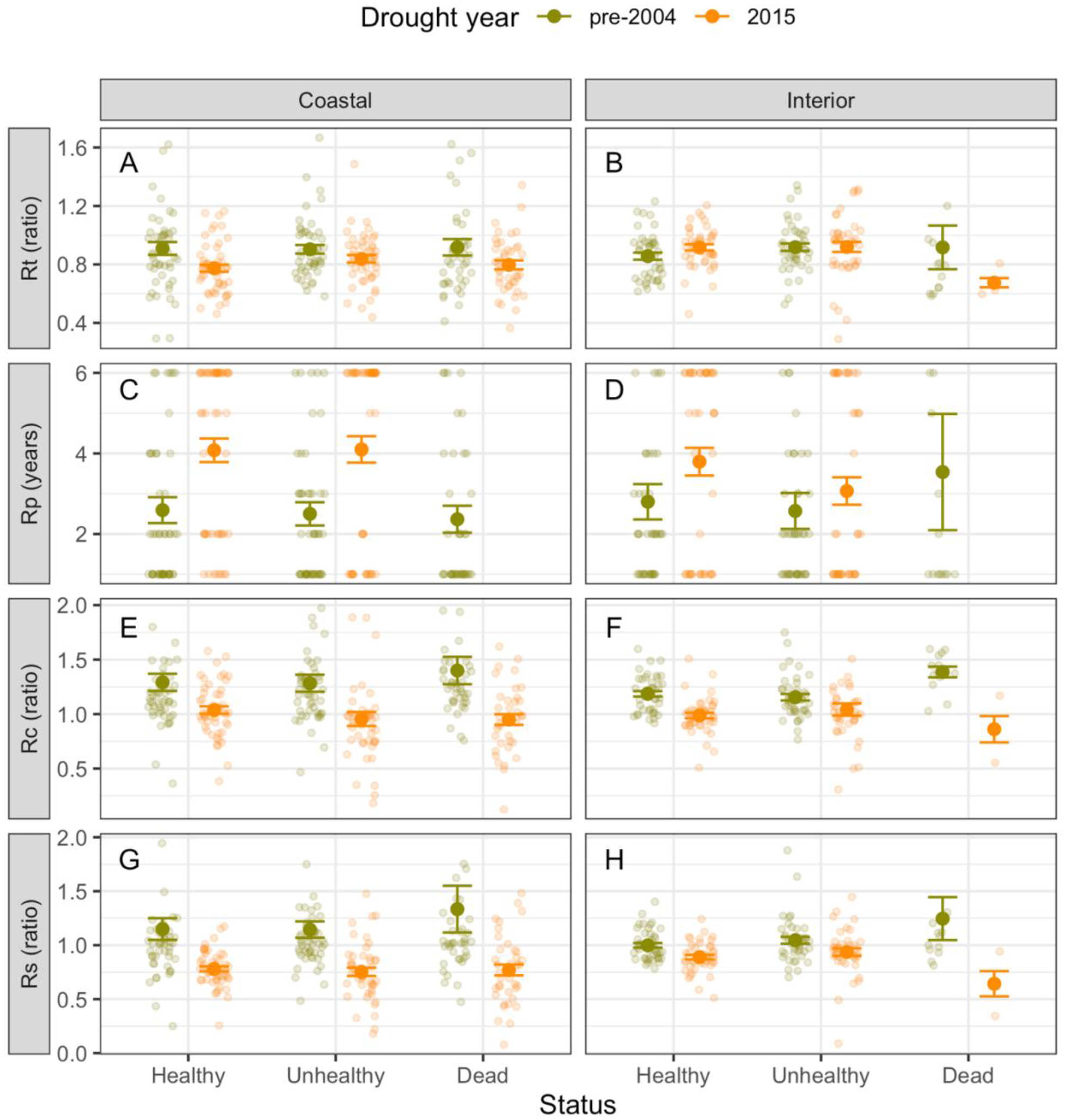
Mean (SE) of four resilience indices characterizing the response of radial growth to two drought events (green, pre-2004 drought; orange, 2015 drought) by tree status for coastal and interior populations of western redcedar (see Table 3 for statistically significant differences among groups). The resistance index (Rt) is the annual growth during the drought year divided by the average of growth in the three years pre-drought. The recovery period (Rp) is the number of years necessary post-drought to return to the average of growth in the three years pre-drought. The recovery index (Rc) is the average of growth three years post-drought divided by annual growth during the drought year. The resilience index (Rs) is the average of growth in the three years pre-drought divided by the average of growth in the three years post-drought. The 2015 drought was followed by much warmer and drier climate conditions than pre-2004 drought events (Fig. 5; Appendix S1: Fig S12).

The pre-2004 drought years were followed by cooler/wetter climate conditions, whereas the 2015 drought year was followed by relatively warmer/drier conditions in interior and coastal populations (Fig. S9). Following the 2015 drought, 77% (110 of 142) and 64% (65 of 101) of trees in coastal and interior populations, respectively, had not returned to pre-drought growth rates (average of prior three years) when late spring and summer SPEI values were again less than -1.0 in 2017 and/or 2018 (Fig. S8). Five years after the 2015 drought year in 2020 (last full ring), growth rates of 50% (50 of 101) and 64% (57 of 88) of healthy and unhealthy trees, respectively, were still below pre-drought growth rates. Consequently, the time required to return to pre-drought growth rates (i.e., recovery period) was significantly longer following the 2015 drought year (mean 3.6 years, SE 0.2) compared to the pre-2004 drought years (mean 2.6 years, SE 0.2) for interior and especially coastal populations, a finding confirmed by the Rp LMM (‘Drought (2015)’ variable; Table 3, Fig. 4C-D).

Recovery of growth rates post-drought relative to the drought year (i.e., recovery index) and the pre-drought period (i.e., resilience index) were significantly slower following the 2015 drought compared to pre-2004 drought years (‘Drought (2015)’ variable in Rc and Rs LMMs, Table 3, Fig. 4E-H), with some important differences by tree status and population (Table 3).

Trees that died after 2015 recovered more rapidly from the pre-2004 drought years and more slowly from the 2015 drought year compared to unhealthy and healthy trees, growth rates of (‘Drought (2015)*Status (dead)’ variable in Rc LMM; Table 3). Additionally, growth rates of trees in interior populations were more resilient to droughts than those in coastal populations, especially following the 2015 drought year (‘Population (interior)’ and ‘Population (interior)* Drought (2015)’ variables in Rs LMM, Table 3).

### Effects of climate variability on tree mortality (Question 4)

In coastal populations, 80% of sampled WRC tree mortality occurred in 2017 or 2018 (as estimated by last partial tree ring), but mortality occurred in all years from 2015 to 2020 (Fig. 5D). Years with WRC tree mortality in coastal populations occurred during exceptionally warm and dry climate conditions as indicated by May to September SPEI of -1.5 in 2015, -0.6 in 2016, -1.0 in 2017 and -1.8 in 2018 (Fig. 5A) and confirmed by daily weather conditions (Fig. 5B-C). From 1970 to 2020, the MTI indicated many hot summer days from 2015 to 2018, with MTI greatest in 2015 and 2018 (Fig. 3B). During the same period, DPI indicated the longest dry periods in 2017 (Everett, WA) or 2018 (Portland, OR; Fig. 5C). Thus, mortality in coastal populations occurred during the same years as exceptionally hot and dry summer conditions (2017, 2018) which was preceded by exceptionally warm and drier conditions in 2015 (i.e., triple whammy). During the years with WRC tree mortality, October to September SPEI was average to cooler and wetter (mean 0.46, range -0.68-1.31; Fig. S10).

**Fig. 5:**
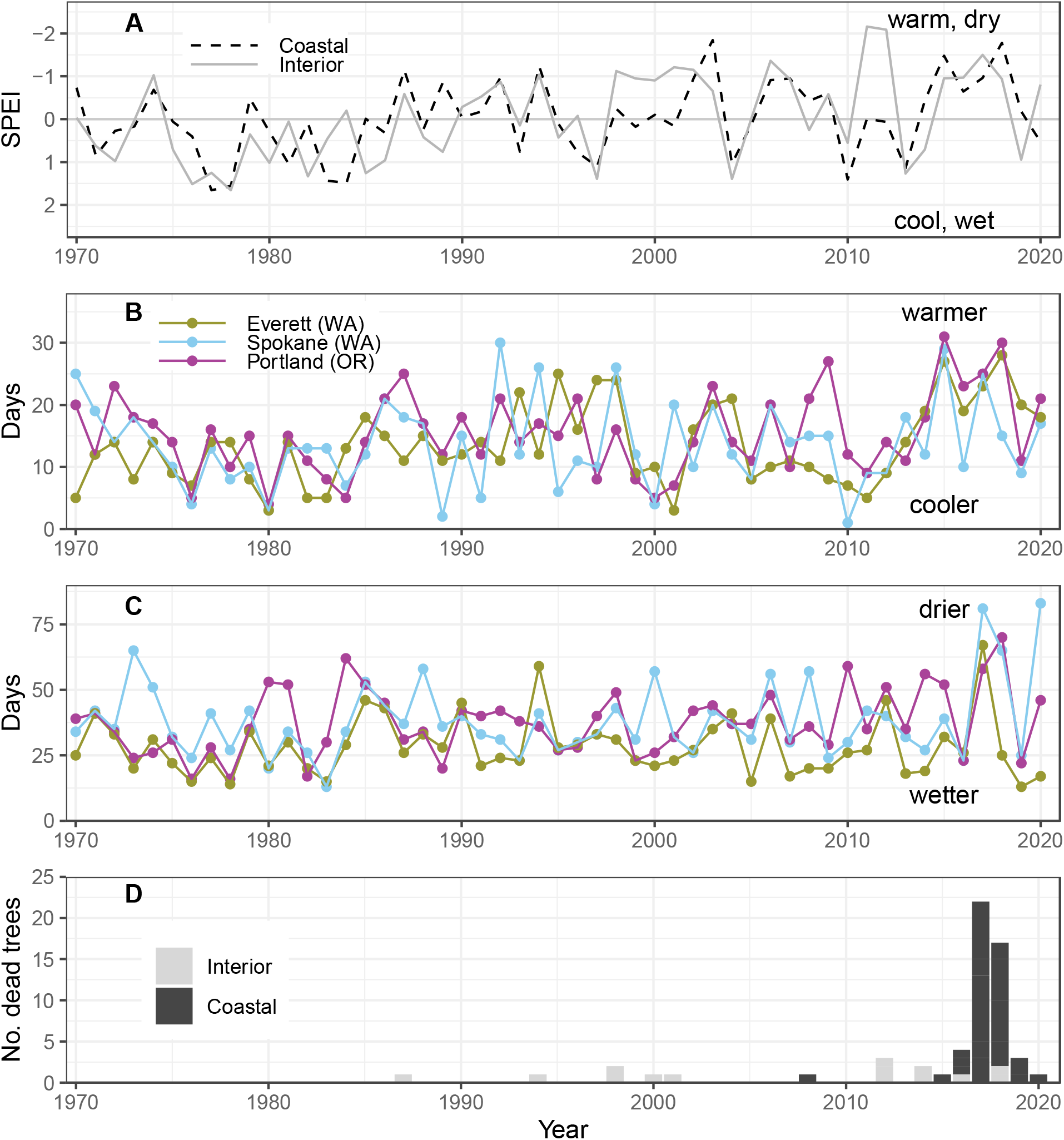
**A)** Standardized Precipitation Evaporation Index (SPEI) averaged for individual years by sites in interior (dashed black, August-September SPEI) and coastal (solid black May-September SPEI) western redcedar populations from 1970 to 2020 (y-axis is inverted). **B)** The Maximum temperature index (MTI) or the number of days maximum temperature exceeded the 90^th^ percentile (May-September) at three weather stations with long-term climate data near the study sites (Fig. 1). **C)** The dry period index (DPI) or the longest consecutive period of days with < 1.0 mm of precipitation (May-September). **D)** The number of dead trees by site and year from 1970-2020.

In interior populations, intermittent, low-level mortality (1-3 sampled trees per year) was reconstructed since the late 1980s, but the majority of sampled WRC tree mortality occurred from 2012 to 2018 (57%; (Fig. 5D). Years with WRC tree mortality were warmer/drier from August to September as evidenced by SPEI < -0.9 for 7 of 9 mortality years (Fig. 5, Fig. S10).

Daily weather conditions indicated that most mortality years (7 of 9) coincided with warmer and/or drier than average conditions from May to September (Fig. 5B-C), as evidence by the MTI (mean 13 days) and DPI (mean 38 days) above their respective mean from 1970 to 2020 (Fig. 5B-C). October to September SPEI during the years with mortality were warmer/drier (4 of 9; Fig. S10), but also cooler/wetter (5 of 9).

## DISCUSSION

Our results support the expectation that WRC tree canopy dieback and mortality in coastal and interior populations were linked to warmer and drier summer climate conditions (Fig. 5). Evidence from tree rings corroborated the reported decline in canopy condition of WRC trees and indicated a period of reduced radial growth prior to mortality that varied by population (Fig. 2). By testing radial growth responses to interannual climate variability (Fig. 3) and drought events (Fig. 4), we identified climate conditions that may increase tree stress and reduce the chances for survival of WRC trees. Warmer and drier conditions in late spring and early summer reduced radial growth and slowed recovery from drought events (Fig. 3-4), whereas a triple whammy of anomalously warm, dry summer conditions was associated with all mortality in coastal populations and most mortality in interior populations (Fig. 5). Collectively, our results are supporting evidence for extreme warm and dry weather and climate as the proximate cause of recent canopy dieback and imply that continued warming and drying climate during the summer will have negative consequences for persistence of WRC trees.

As expected, we found that radial growth of dead and unhealthy trees (to a lesser degree) diverged from healthy trees prior to death or 2020, respectively, in coastal and interior populations of WRC trees (Fig. 2). Tree death was portended by an average of 4-5 years of radial growth decline that culminated in the lowest growth the year prior to death relative to healthy trees, a new finding for WRC trees. Similar reductions in radial growth were also observed for many other gymnosperms prior to death, including Cupressaceae Alaskan cedar (*Callitropsis nootkatensis*; (Comeau et al. 2019)), though the period and magnitude of decline varies widely by species (Cailleret et al. 2017). Reductions in tree growth can be a key indicator of declining tree vigor and a higher likelihood for mortality (Bigler et al. 2004). However, WRC trees with partial canopies and trees that died endured multiple decades to just a few years of reduced tree growth (relative to comparable surviving trees). Thus, reduced WRC tree growth does not necessarily indicate imminent mortality and variability in the timing of the growth reduction does not help identify climate in a specific year in the last decade as the ‘inciting’ event for mortality during the 2015-2019 period. However, leading up to 2015, the first of four exceptionally warm/dry years, many of the trees that died in 2017 or 2018 were growing slower than neighboring healthy trees. Assessment of the interannual variability in growth prior to death, coupled with eco-physiological measurements, for trees that survived and died may produce more accurate predictors of future tree mortality (Cailleret et al. 2019).

In our study, drier and warmer climate conditions in May and/or June reduced radial growth of WRC trees in many coastal and interior sites (9 of 11 sites; Fig. 3). Tree health status did not affect these climate-growth relationships, potentially due to the relatively short period (∼5 years) in which radial growth of dead and unhealthy trees differed from healthy trees (i.e., compared to the longer 1975-2020 analyzed period). Consistent with our results on climate-growth relationships, height growth of WRC trees occurred from mid-May to mid-July (data from 1961; (Walters and Soos 1963)) and radial growth decreased during warmer June maximum temperatures and lower April to July total precipitation than average in dry sites in coastal populations of WRC trees in southern British Columbia, Canada (Seebacher 2007). The amount of precipitation and the temperatures in May and June likely regulates the availability of soil moisture for photosynthesis, carbon gain, and radial growth during the growing period (mid-May to mid-July) and the rate that soil moisture is depleted during the subsequent and consistently dry period from July to September (Baker et al. 2019). Greater precipitation and cooler temperatures in May and June may prolong the period of ample soil moisture for tree growth further into the summer months, whereas warm, dry climate conditions in May and June may increase the length of the summer soil moisture drought, potentially increasing tree stress and susceptibility to canopy dieback and mortality. In summary, our finding for WRC trees, that growth is limited by late spring and early summer moisture availability, is consistent with multiple other conifer species in the PNW region, such as western hemlock (Brubaker 1980, Nakawatase and Peterson 2006).

Warmer and drier late spring and summer conditions also slowed recovery of radial growth following droughts in our study (Table 3; Fig. 4). Most trees (>65%) were still in the process of recovering radial growth from the 2015 drought when two consecutive years of warmer, drier late spring and summer climate conditions occurred in 2017 and 2018.

Consequently, radial growth rates required longer to recover (Rp) and were notably less resilient (Rs) following the 2015 drought compared to the pre-2004 drought years with cooler, wetter post-drought conditions (Table 3). WRC trees close their stomata to limit water loss during periods of low soil moisture availability (Grossnickle et al. 2005) and are more vulnerable to cavitation for some organs and can have smaller root systems than co-occurring species (e.g., Douglas-fir) (Mcculloh et al. 2014). Warm and dry conditions during the recovery period may have slowed recovery of water transport capacity (due to cavitation) and lengthened the period of stomatal closure, thereby lowering photosynthesis and reducing carbon gain, as has been observed for other conifers (Brodribb and Cochard 2009, Trugman et al. 2018).

Our findings support the hypothesis that *naïve* populations exposed to moderate climate conditions (coastal) are more susceptible to changes in climate than those populations adapted to more variable, continental climate (interior; Bonebrake and Mastrandrea 2010). Compared to coastal populations, radial growth rates of WRC trees in interior populations were more resistant and resilient to drought (Fig. 4, Table 3), though drought severity was slightly lower at sites in interior populations in 2015 (Fig. S9). The higher growth resistance of WRC trees in interior relative to coastal populations is consistent with their greater water use efficiency and drought tolerance, specifically osmotic potential and relative water content at turgor loss point (Grossnickle and Russell 2010). Though there is minimal genetic variation north to south within WRC populations (Shalev et al. 2022, Rehfeldt 1994), differences between populations in physiological and growth response to drought in the southern half (climatically warmer and drier) of its range may indicate that interior populations are better adapted to continued warming and drying summer conditions.

Spatial and temporal synchrony in unfavorable climate conditions and tree mortality (e.g., over ∼400km in 2017 and 2018 in coastal populations) is strong evidence that climate is a key contributor to WRC tree mortality. Consistent with radial growth responses to climate variability and drought, we found that WRC tree mortality was associated with warm and dry summer conditions in coastal and interior populations (Fig. 5). However, the timing and quantity of mortality and the specific climate conditions associated with mortality differed between populations. WRC tree mortality in sampled coastal populations was episodic (80% of dead trees sampled had mortality in two years, 2017-2018), whereas mortality in interior populations occurred over a longer period with fewer deaths of sampled trees per year (Fig. 5). In coastal populations, WRC tree mortality was associated with anomalously warm and dry late spring and summer climate conditions from 2015 to 2018–including exceedingly high maximum summer temperatures and the longest summer dry period from 1970 to 2020 (i.e., hot drought). The WRC trees that died in the two to three years following the 2015 drought year recovered more rapidly to prior droughts of similar severity (i.e., higher resilience) and had a similar resistance to the 2015 drought as healthy and unhealthy trees. However, the trees that died lacked the capacity to recover to subsequent droughts during the recovery period (i.e., compounded drought; Peltier et al. 2016). Similar repeated years with warm and dry conditions may not have occurred since the 1930s (as indicated by SPEI, Fig. S8), and nearly all dead WRC trees that we sampled in coastal populations likely germinated after the 1940s (Fig. S1) or were shaded by overstory trees during the 1930s warm and dry period. In interior populations, WRC tree mortality was limited but associated with warm and dry conditions in late summer (August to September). Climate conditions associated with mortality in both populations would have likely resulted in low soil moisture availability, high evaporative demand, and a greater chance for hydraulic failure (McDowell et al. 2022).

By investigating the varied responses of WRC tree growth and canopy dieback to climate variability, our findings prove insightful for understanding the implications of climate change.

Our results suggest that seasonal rather than annual shifts in temperature and precipitation may have the greatest effect on WRC tree growth and dieback, though we were not able to evaluate the effect of long-term increases in air temperature. Based on the positive correlations between maximum temperature and radial growth from November to January in interior populations, we infer that continued warming in fall and winter months may increase tree growth (Kunkel et al. 2013), though the mechanism for this correlation needs exploration. In contrast, projected decreases in summer precipitation (Mote and Salathé 2010) would reduce growth of WRC trees in lower elevations in both populations (present study) and in high elevation habitats in the Cascade Mountains (Ettinger et al. 2011).

Of greatest concern for the persistence of WRC trees and possibly other co-occuring species is the increasing trend in growing season climate water deficits, decreasing trend in summer soil moisture availability, and the forecasted increase of nine days in the annual maximum consecutive days with < 3 mm of precipitation by mid-century in the PNW region (Kunkel et al. 2013, Gergel et al. 2017). Such conditions were associated with recent WRC tree dieback in lower elevation habitats in interior and coastal population (Fig. 5). Although not addressed in our study, mortality and dieback of three common associates of WRC trees (western hemlock, grand fir, and bigleaf maple) in the last decade and over a smaller area than WRC dieback may be related to unfavorable climate conditions (WA DNR 2020, Betzen et al. 2021).

For WRC trees, our results imply that rapid shifts toward warmer, drier summer climate conditions would further reduce tree growth and elevate the risk of canopy dieback and possibly other co-occurring species and species with similarly low drought tolerance in the Cupressoid clade (Cupressaceae family; Pittermann et al. 2012).

Our research needs to be considered in the context of where we sampled. In coastal populations, recent WRC dieback was observed and sampled in low elevation, warmer and drier forests with relatively young WRC trees (<150 years; except IC site). We explored higher elevation WRC forests (Cascade and Olympic Mountains) and lower elevation forests in wetter locations (temperate rainforests along Washington and Oregon coast) and found evidence of old, partial canopy dieback (e.g., spike tops). However, we did not observe mortality from the last two decades, and thus did not sample these locations. In interior populations, sampling included lower and higher elevation sites with younger (<150 years) and older WRC trees (>150 years), but a relatively small sample size of dead trees. Future research is needed that explores how tree- (e.g., tree size, age, and height as well as root pathogens) and stand-scale factors (e.g., tree density, soils, topography, and location within the distribution of WRC) may increase vulnerability to climate-caused dieback (Hennon et al. 2020).

## CONCLUSION

By exploring WRC as a ‘canary in the forest’, our findings are an early warning that warming climate and multi-year hot droughts during the late spring and summer will likely increase the vulnerability of trees to canopy dieback. Our results provide key context for managing a culturally irreplicable species under changing climates. Global syntheses of tree mortality indicate no or very few tree mortality events from hot droughts in the PNW, and we show a direct effect of climate and weather on tree mortality in low elevation, mesic forests in the PNW region in the absence of fire or primary biotic mortality agents. Continued loss of WRC trees may shift tree species composition to species more tolerant of current and future climate conditions (e.g., Douglas fir). Depending on the quantity of mortality, reduced stand density may increase resource availability for survivors, potentially supporting increases in tree vigor and growth, with responses likely varying by site conditions and population. In the context of adapting forests to future climate conditions, WRC trees in the interior population appear more drought-resistant and less vulnerable to widespread, episodic canopy dieback than coastal populations. Selecting seeds from provenances or genotypes in interior populations may help sustain WRC trees in coastal populations and assist with migrating WRC trees to match the rapid changes in global climate.

## ACKNOWLEDGEMENTS

For field assistance, we thank Isaac Ball and Tiago Holz. For helpful feedback on earlier versions of this manuscript, we thank Constance Harrington and Dave L. Peterson. This research was funded by the US Forest Service Region 6 Emerging Pest Investigation (Agreement #21-CA-11062765-742), McIntire Stennis project (WNP00009), and USDA NIFA postdoctoral award (2022-67012-37200).

## DATA AVAILABILITY

The data that support the findings of this study are available from the corresponding author upon reasonable request.

## STATEMENT OF AUTHORSHIP

RAA, AH, AJHM, KBM, AR, CJB, MF, BAG and HDA designed the study. RAA, HDA, BM, MF, BAG, and AH collected the tree cores. RAA, LRP, BM, JTY, and ARC performed laboratory work. RAA performed the statistical analysis and wrote the initial draft of the manuscript. All authors contributed to writing and revising the manuscript.

## CONFLICT OF INTEREST

The authors declare that they have no competing interests.

## Notes

### Competing Interest Statement

The authors have declared no competing interest.

