## Supplementary Materials for "Canary in the Forest? – Tree mortality and canopy dieback of western redcedar linked to drier and warmer summer conditions"

### Supplementary Materials Table of Contents

**Table S1:** Abiotic and biotic characteristics of the sampled stands

**Table S2:** Climate characteristics of the sampled stands

**Fig. S1:** Sample depth of tree ring chronologies by year for all, healthy, unhealthy, or dead trees by site

**Fig. S2:** Western redcedar tree ring chronologies

**Fig. S3:** Correlation coefficients between total monthly precipitation and the site chronology by site

**Fig. S4:** Correlation coefficients between monthly maximum temperature and the site chronology by site

**Fig. S5:** Correlation coefficients between Standardized Precipitation Evaporation Index (SPEI) and the site chronology by site

**Fig. S6:** Average count of statistically significant climate-growth relationships by tree status and period

**Fig. S7:** Average correlation coefficients from climate-growth relationships by tree status and period

**Fig. S8:** May to June and May to September Standardized Precipitation Evaporation Index (1900-2020)

**Fig. S9:** SPEI pre-drought, during the drought year, and post-drought for the pre-2004 and 2015 drought years

**Fig. S10:** Scatter plots of SPEI during years with and without WRC tree mortality

**Fig. S11:** Average basal area increment (BAI) by site by tree status and population (1975-2020)

**Fig. S12:** Annual growth ratios for individual trees by population.

**Fig. S13:** Correlations table of site tree ring chronologies by tree status (1975-2020)

**Table S1:** Abiotic and biotic characteristics of the 11 sampled stands with western redcedar (WRC, including location (decimal degrees), slope, aspect, and description of the stand. Stand descriptions based on field notes. See Table 1 for sample depth and other characteristics

| Pop. | Site | Lat. | Long. | Slp (˚) | Asp. (˚) | Stand description |
|---|---|---|---|---|---|---|
| Interior | RG | 48.7676 | -117.0609 | <5 | flat | Monotypic stands of western redcedar intermixed with mesic mixed conifer forest with western redcedar, grand fir, western hemlock, and western larch. No evidence of logging and few thinning trees. |
| | HG | 47.0869 | -116.1183 | 5-10 | 340 | Monotypic stand of WRC (~ 20ha) with a few grand fir. No evidence of logging but surrounded by clear-cuts. |
| | SL | 48.9712 | -117.3332 | <5 | 240 | Post-fire stand (fire scars) and no evidence of logging with mesic mixed conifer species composition, including Douglas fir, western larch, western redcedar, and western white pine. |
| | MP | 47.0650 | -116.8985 | 10-15 | 30 | WRC dominated stand (~80% by basal area) with grand fir and western larch. Likely a post-fire stand as evidence by fire scars and rotting paper birch on the forest floor. No evidence of logging. |
| | LI | 48.5100 | -117.9487 | <5 | 75 | WRC dominated stand (~85% by basal area) located adjacent to ephemeral wetland. Last harvest 1968 (WA Dept. of Natural Resources; no additional details). |
| Coastal | BW | 45.3842 | -122.0396 | <5 | flat | Monotypic stand of WRC trees on riverbed terrace adjacent to Sandy River. No evidence of flood damage or logging (cut stumps). |
| | LM | 48.3986 | -122.3019 | 10-15 | 75 | Mesic mixed species composition with western redcedar, bigleaf maple, Douglas fir, grand fir. Some logging in the area in early 1990s. |
| | IC | 47.6102 | -120.9450 | <5 | 30 | Mesic mixed conifer species composition with western redcedar, western hemlock, Douglas fir, pacific yew. No evidence of logging. |
| | TR | 47.9895 | -122.7930 | <5 | 60 | Mesic mixed species composition with western hemlock, Douglas fir, western redcedar, bigleaf maple. Last harvest ~1940s. |
| | RF | 45.3095 | -122.6360 | 10-15 | 170 | Mixed species composition with western redcedar, Oregon white oak, bigleaf maple, and Douglas fir. No evidence of logging. |
| | MR | 45.4284 | -123.2311 | <5 | 80 | Mesic mixed species composition with western redcedar, bigleaf maple, Douglas fir. Minimal forest management prior to removal of recently dead WRC trees. |

Site names abbreviations: Roosevelt Grove (RG), Hobo Grove (HG), Slate (SL), McCroskey Park (MP), Lindsey (LI), Barlow Wayside (BW), Little Mountain (LM), Icicle Creek (IC), Trillium (TR), Ramirez Farm (RF), Mount Richmond (MR).

**Table S2:** Climate characteristics of the 11 sampled stands with western redcedar from 1975 to 2020, including mean annual precipitation (MAP), growing season precipitation (GSP, April-September), mean maximum temperature of the hottest month, and minimum temperature of the coldest month. Data from PRISM (2022).

| Pop. | Site | MAP (mm) | GSP (mm) | MAT (˚C) | Max. temp. hottest month (˚C) | Min. temp. coldest month (˚C) |
|---|---|---|---|---|---|---|
| Interior | RG | 1073.5 | 283.5 | 4.0 | 28.3 | -17.2 |
| | HG | 1294.5 | 285.1 | 5.4 | 29.0 | -16.2 |
| | SL | 733.6 | 256.6 | 6.4 | 32.1 | -18.6 |
| | MP | 790.8 | 195.7 | 7.1 | 31.1 | -16.9 |
| | LI | 575.9 | 212.1 | 7.6 | 32.7 | -15.5 |
| Coastal | BW | 2350.6 | 471.2 | 9.2 | 27.7 | -6.9 |
| | LM | 1092.5 | 275.3 | 10.1 | 26.6 | -4.4 |
| | IC | 1730.2 | 237.4 | 5.3 | 26.9 | -13.7 |
| | TR | 1249.4 | 160.2 | 11.2 | 30.1 | -4.1 |
| | RF | 1168.2 | 195.8 | 12.2 | 30.7 | -3.3 |
| | MR | 1249.4 | 160.2 | 11.2 | 30.1 | -4.1 |

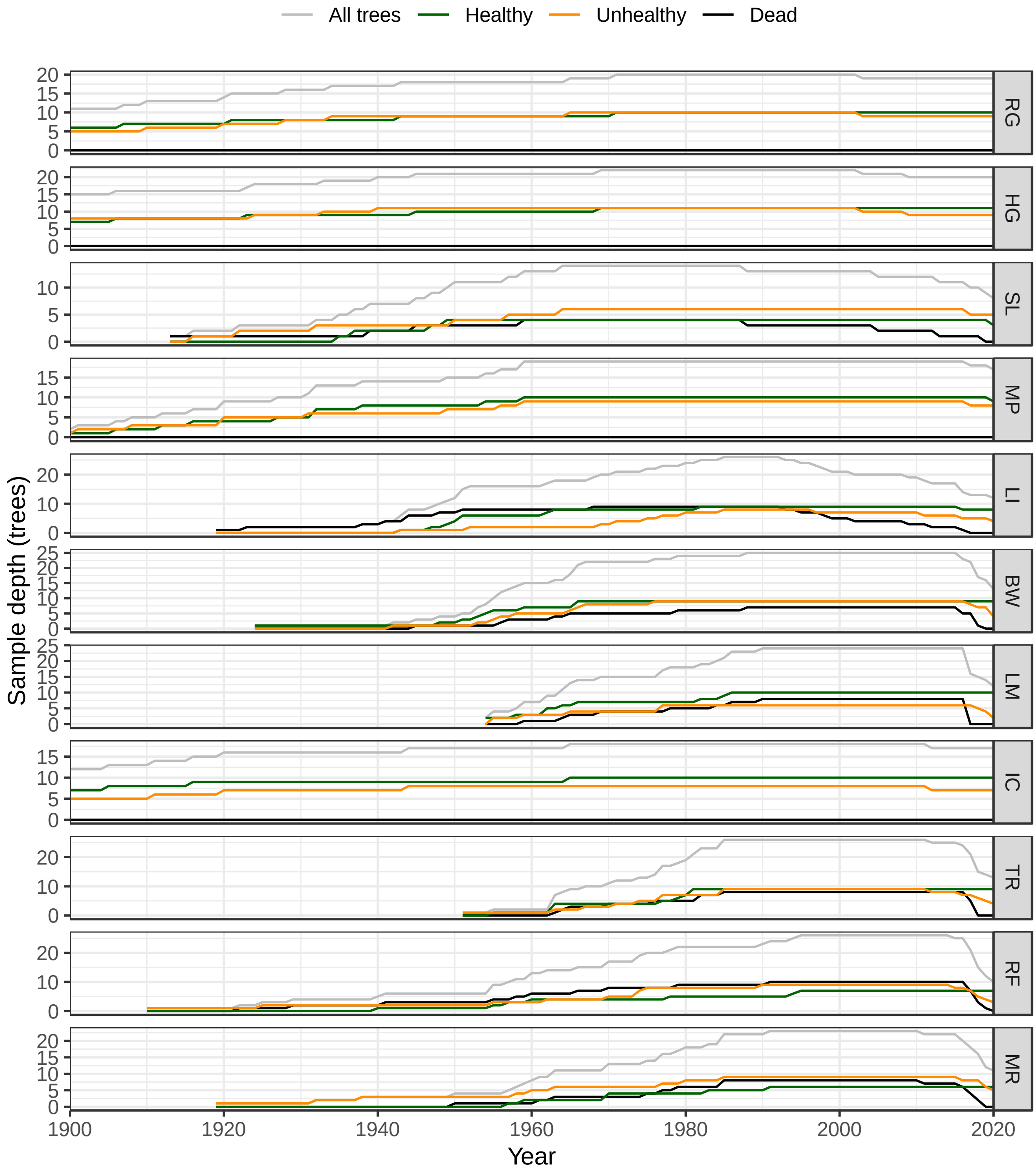


**Fig. S1:** Sample depth of tree ring chronologies by year for all, healthy, unhealthy, or dead trees by site. Note y-axis varies by site.

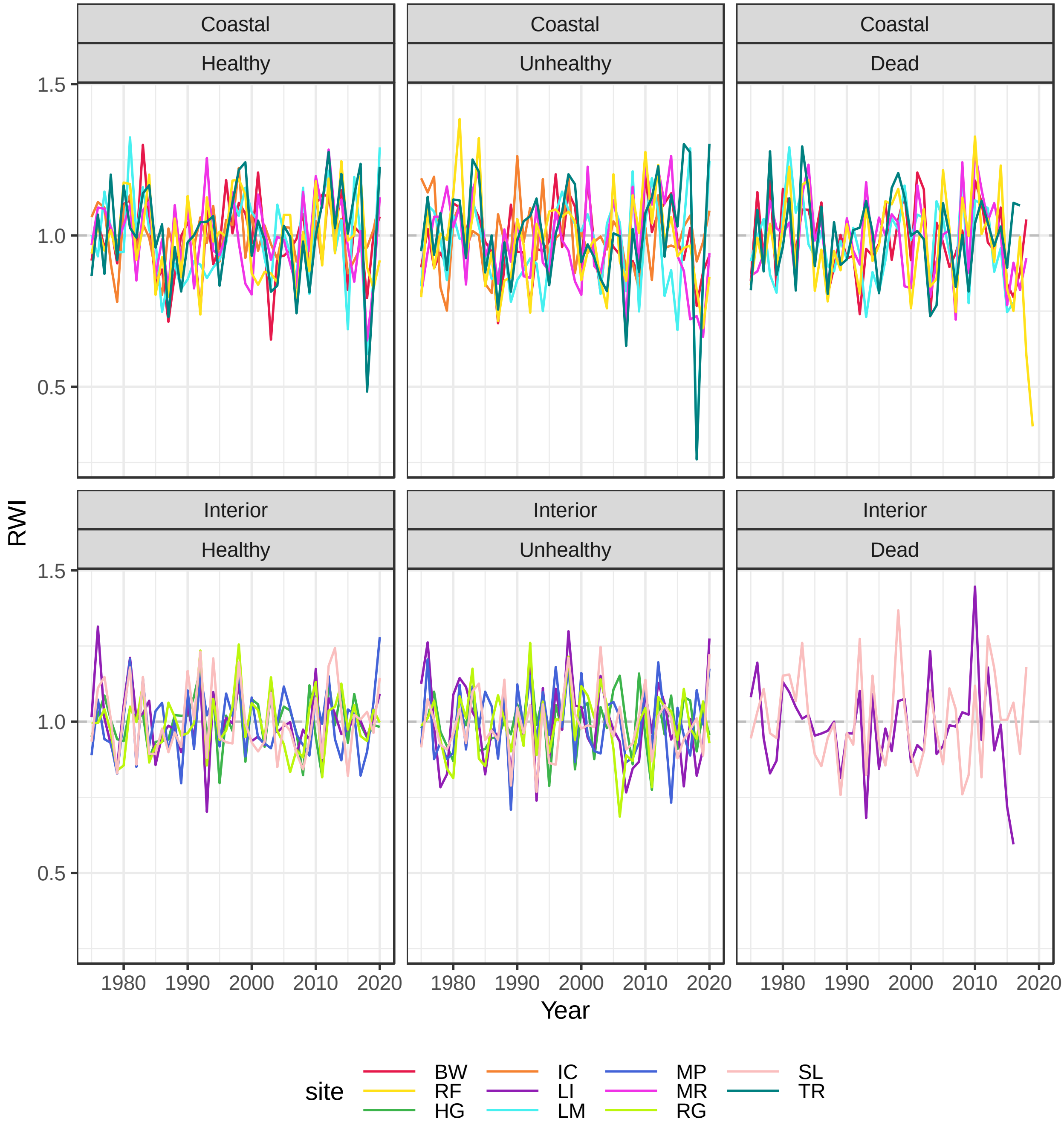


**Fig. S2:** Western redcedar tree ring chronologies, expressed as the ring width index (RWI), by site for healthy, unhealthy (canopy dieback), and trees that died (dead) in coastal and interior populations from 1975 to 2020.

**Tree radial growth response to climate conditions by tree health status and period**

To test whether climate-growth relationships varied by tree status (health, unhealthy, dead) or period of analysis, we tested and then compared correlations between site tree ring chronologies (for each health status group in two periods) and four climate variables: (i) monthly maximum air temperature, (ii) monthly total precipitation, (iii) monthly SPEI, and (iv) vapor pressure deficit ('treeclim' package in R; (Zang and Biondi 2015)). We selected the period 1975 to 2020 for analysis, because each tree health status group chronology had > 5 trees. To test for changes over time in climate-growth relationships, we examined climate-growth relationships from 1975–1999 and 2000–2020 based on the changes in climate around 2000 (Abatzoglou and Williams 2016). We computed correlations for each month from May of year prior to ring formation (lagged effects) to October of ring formation year for each site, which resulted in 1,260 correlation tests per climate variable (11 sites x 2-3 tree health status groups x 18 months x 2 periods; Fig S3-S6 for results). We compared the average count of statistically significant relationships ($\alpha = 0.05$; Fig S7) and the mean of the correlation coefficients by tree status and period for each climate variable and WRC population (Fig. S7). Visual comparison of means and standard errors indicated no significant differences by tree status or period (Fig. S6-S7). To confirm that climate-growth relationships did not vary by tree health status, we tested multiple linear regressions with site chronology by tree status as the response variable and individual monthly climate variables and tree health status as predictor variables from 1975 to 2020. The tree health status predictor variable was not significant ($p > 0.4$). Because the tree health status and period variables had negligible effects on climate-growth relationships, we excluded these variables from analysis in the main manuscript to simplify results.

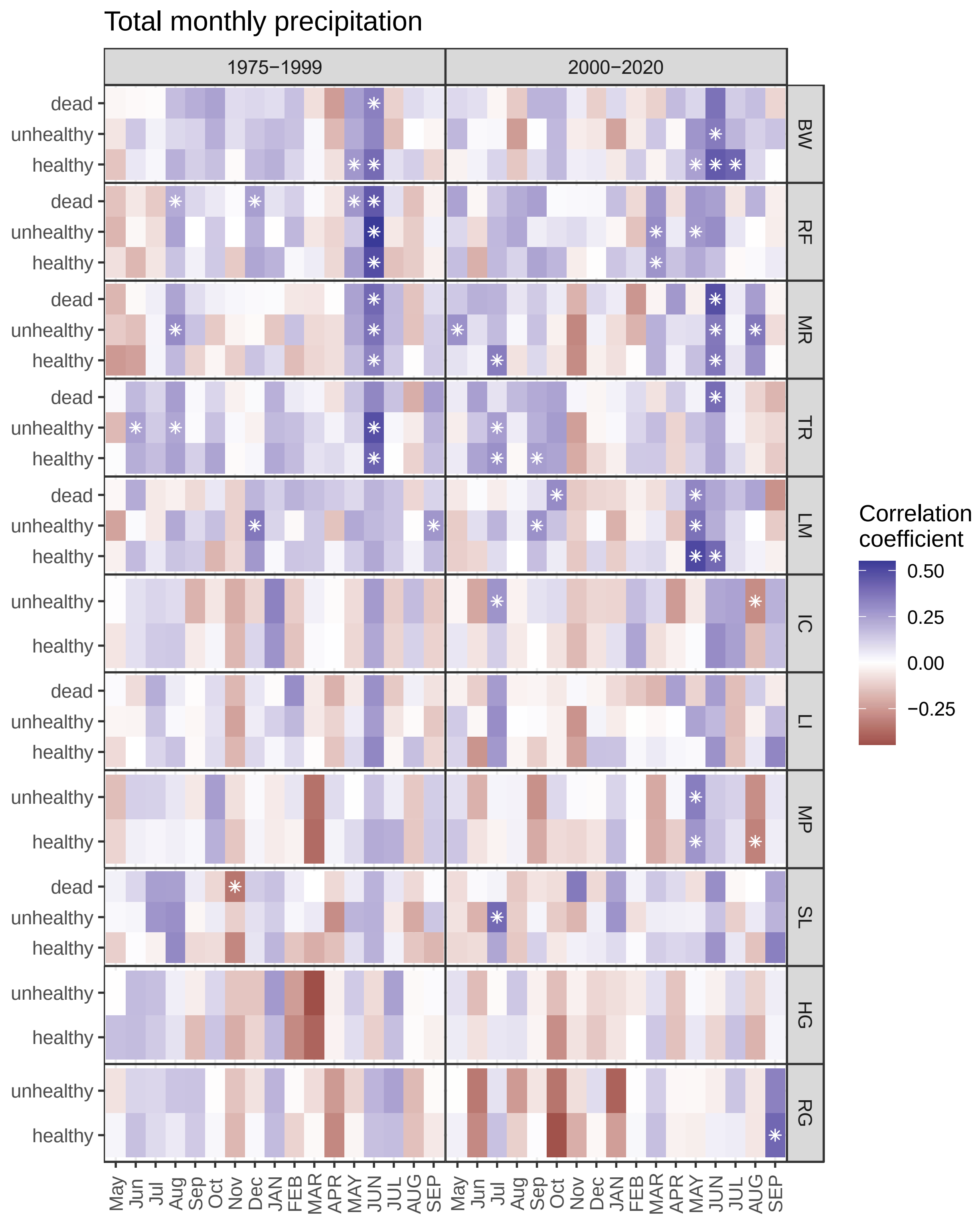


**Fig. S3:** Correlation coefficients between total monthly precipitation and the site chronology by site (right y-axis label) and tree status (left y-axis label) separated by coastal (top rows, BW-IC) and interior (bottom rows, LI-RG) populations of western redcedar during two periods, 1975-2000 (left) and 2000-2020 (right). Monthly climate from the year prior to growth (lower case) to the growth year (upper case) were considered. * $P < 0.05$

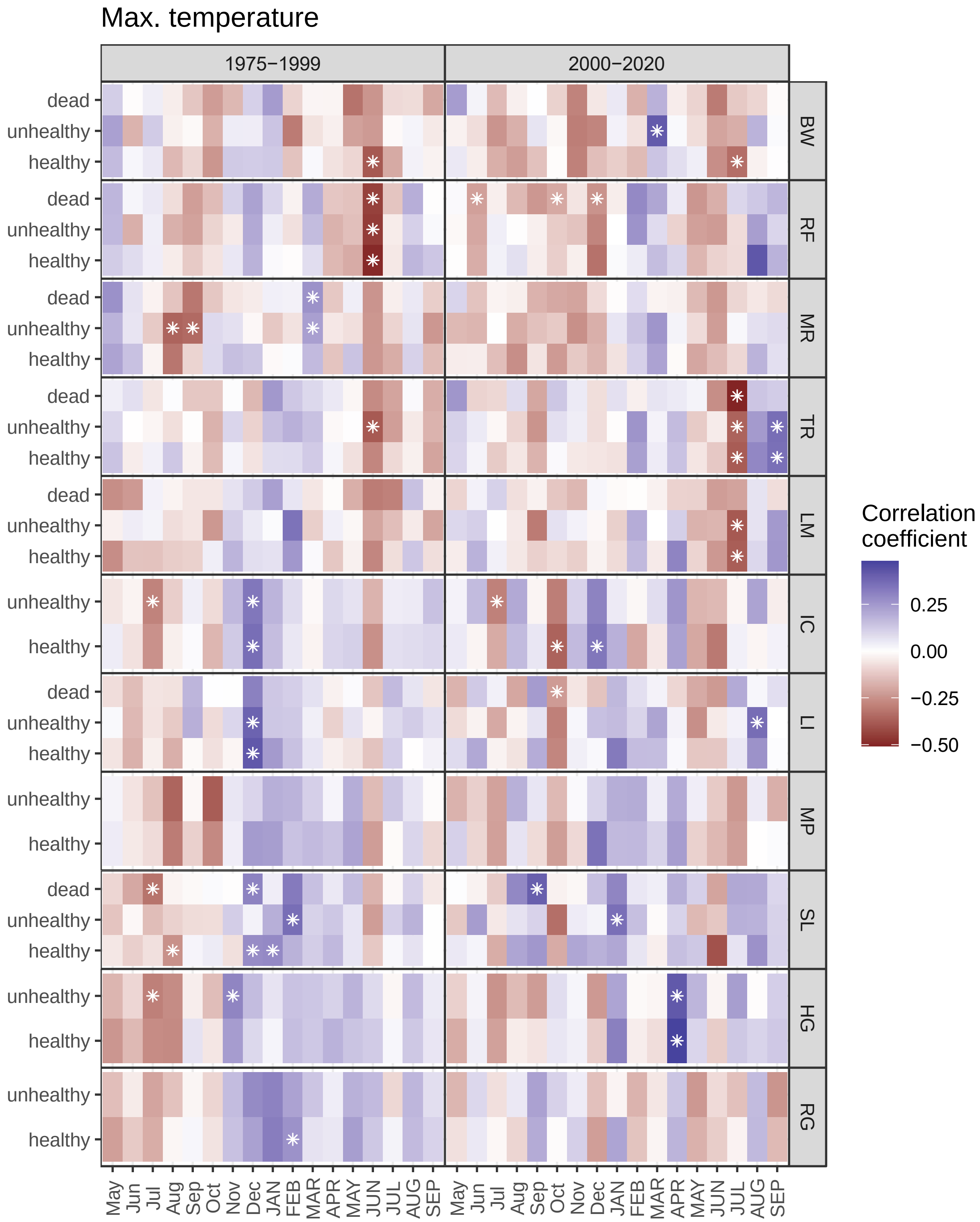


**Fig. S4:** Correlation coefficients between monthly maximum temperature and the site chronology by site (right y-axis label) and tree status (left y-axis label) separated by coastal (top rows, BW-IC) and interior (bottom rows, LI-RG) populations of western redcedar during two periods, 1975-2000 (left) and 2000-2020 (right). Monthly climate from the year prior to growth (lower case) to the growth year (upper case) were considered. * $P < 0.05$

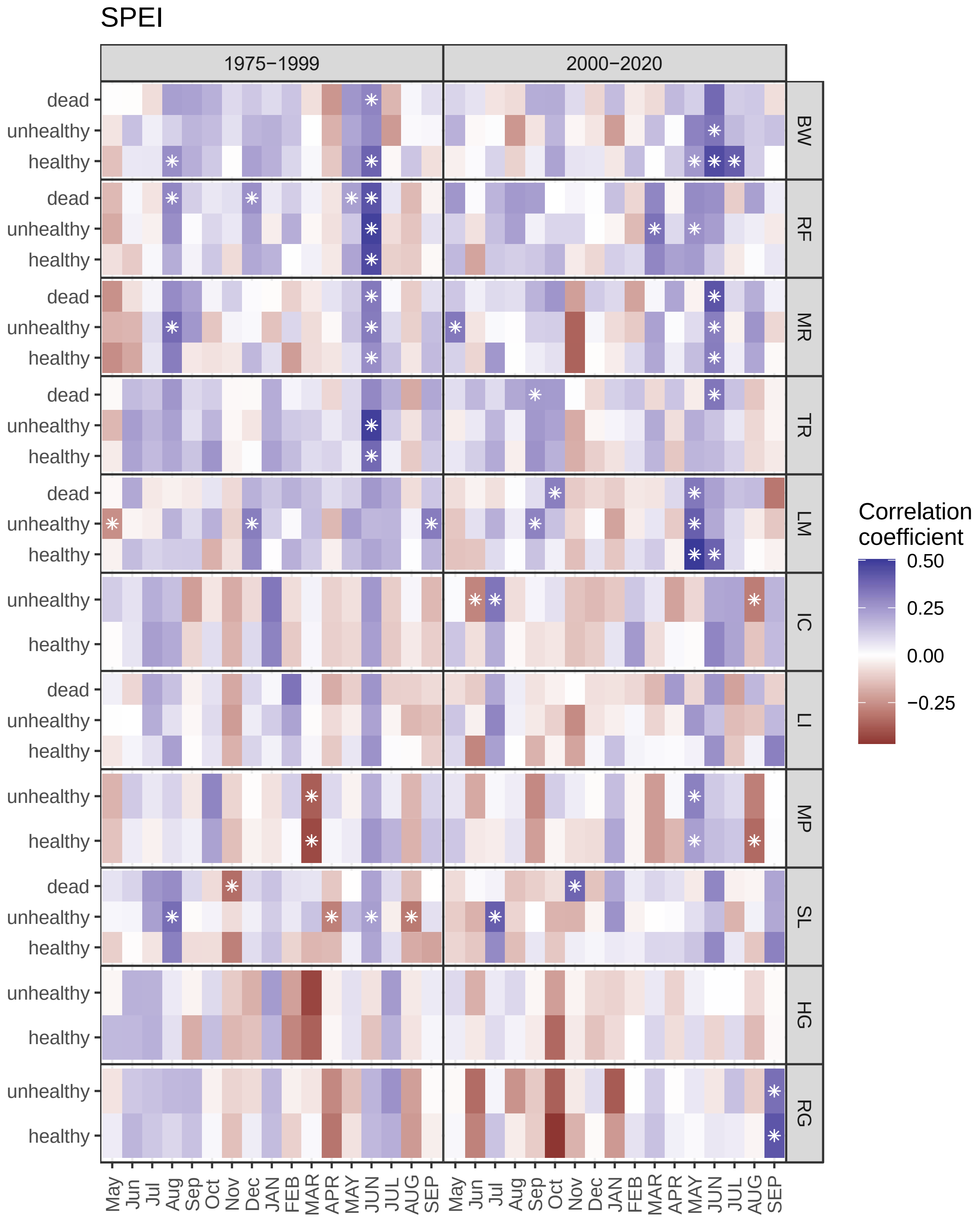


**Fig. S5:** Correlation coefficients between Standardized Precipitation Evaporation Index (SPEI) and the site chronology by site (right y-axis label) and tree status (left y-axis label) separated by coastal (top rows, BW-IC) and interior (bottom rows, LI-RG) populations of western redcedar during two periods, 1975-2000 (left) and 2000-2020 (right). Monthly climate from the year prior to growth (lower case) to the growth year (upper case) were considered. * $P < 0.05$

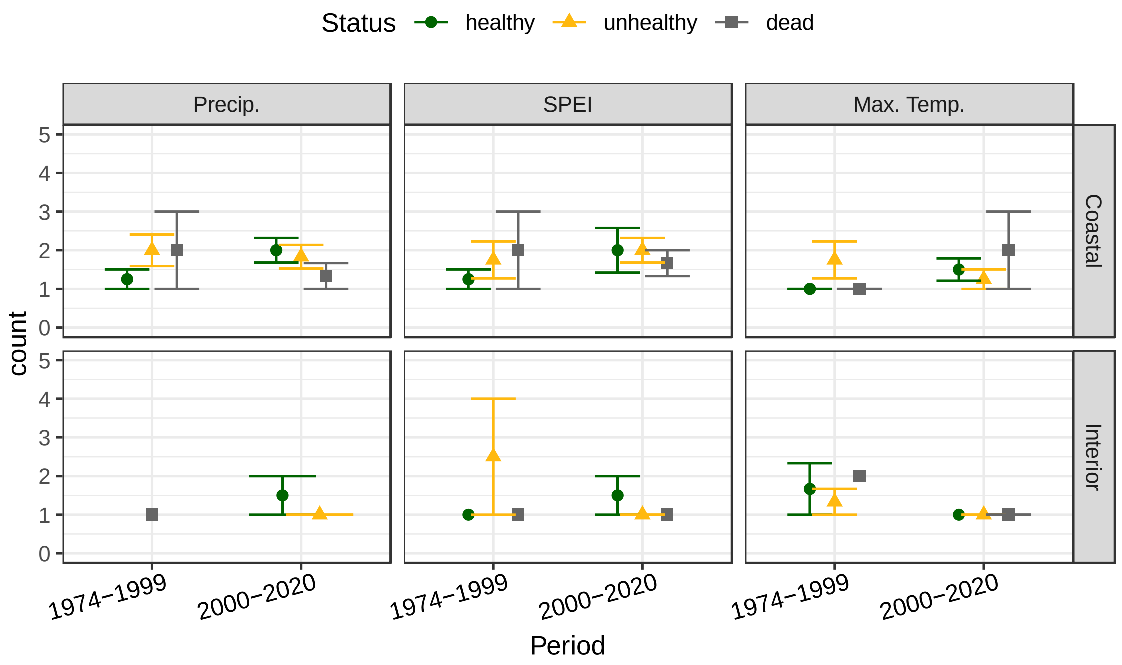


**Fig. S6:** Average count of statistically significant relationships between monthly climate variables and radial tree growth in two periods by climate variable, population, and tree status, summarizing results from Fig. S3-S5. Error bars are the standard error, and points are the site counts.

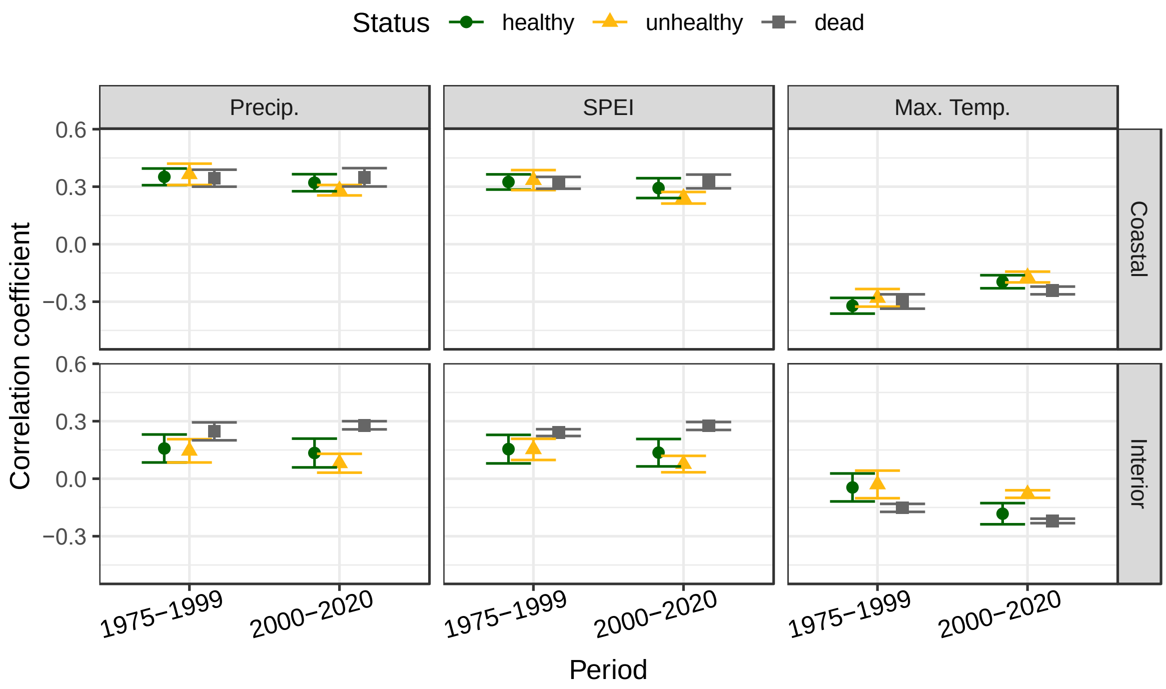


**Fig. S7:** Average correlation coefficients from correlation analysis between climate variables in June of growth year (consistently significant across many sites) and radial tree growth in two periods by climate variable, population, and tree status, summarizing correlation coefficients from Fig. S3-S5. Error bars are the standard error, and points are the site correlation coefficients.

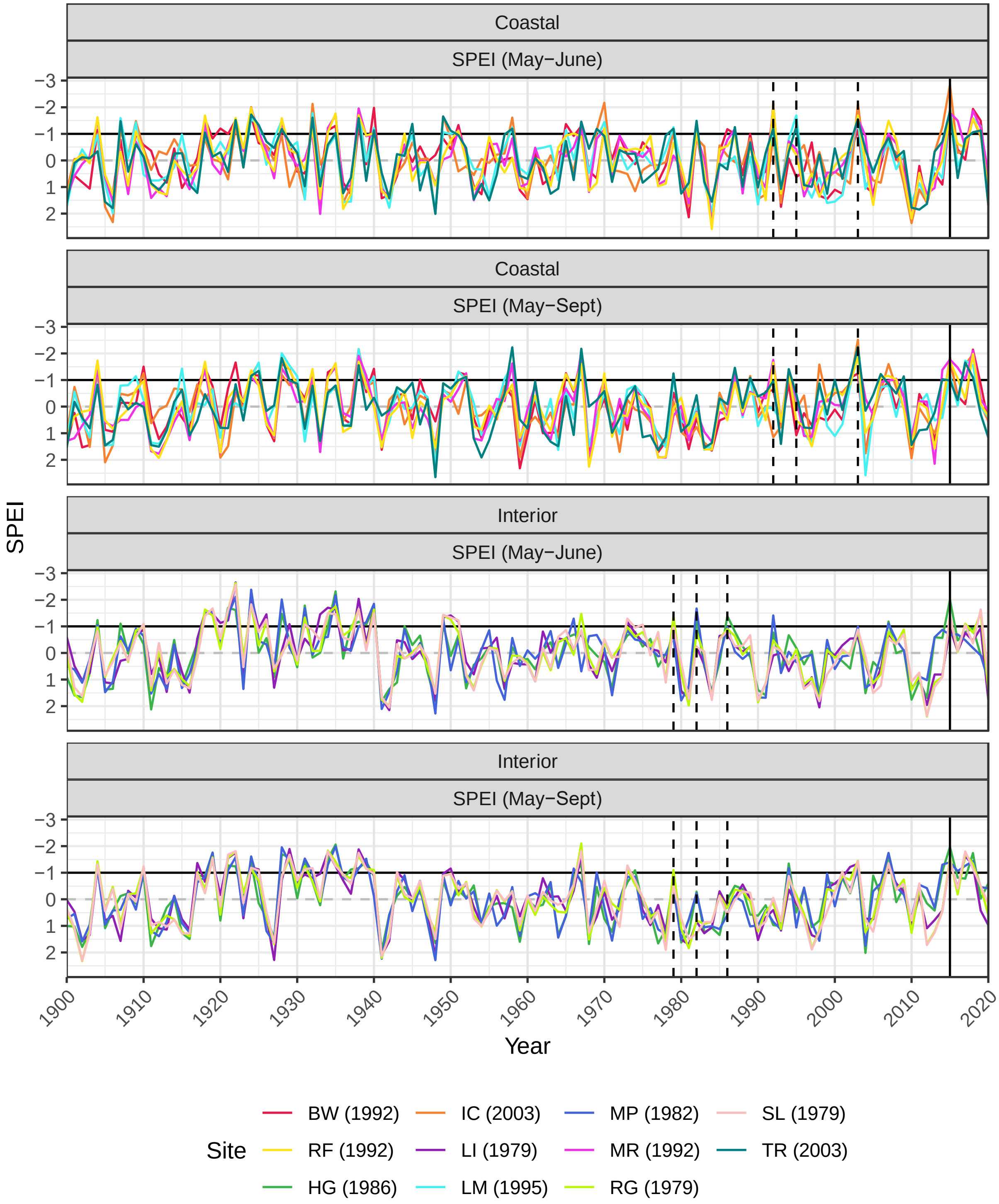


**Fig. S8:** May to June and May to September Standardized Precipitation Evaporation Index (SPEI) for each field site in coastal and interior populations of WRC trees. In our resilience index analysis (Question 3), we selected two exemplary drought years per site with May-June SPEI < -1.0: (i) the 2015 regional drought year for all sites (warm, dry post-drought period; vertical solid line) and (ii) a site-specific drought year (see legend for a site's drought year; dashed vertical line in plot) with the lowest May-June SPEI (other than 2015) and cool, wet post-drought conditions. The graph box indicates the study period, the black horizontal line is at -1.0, and the y-axis is reversely plotted.

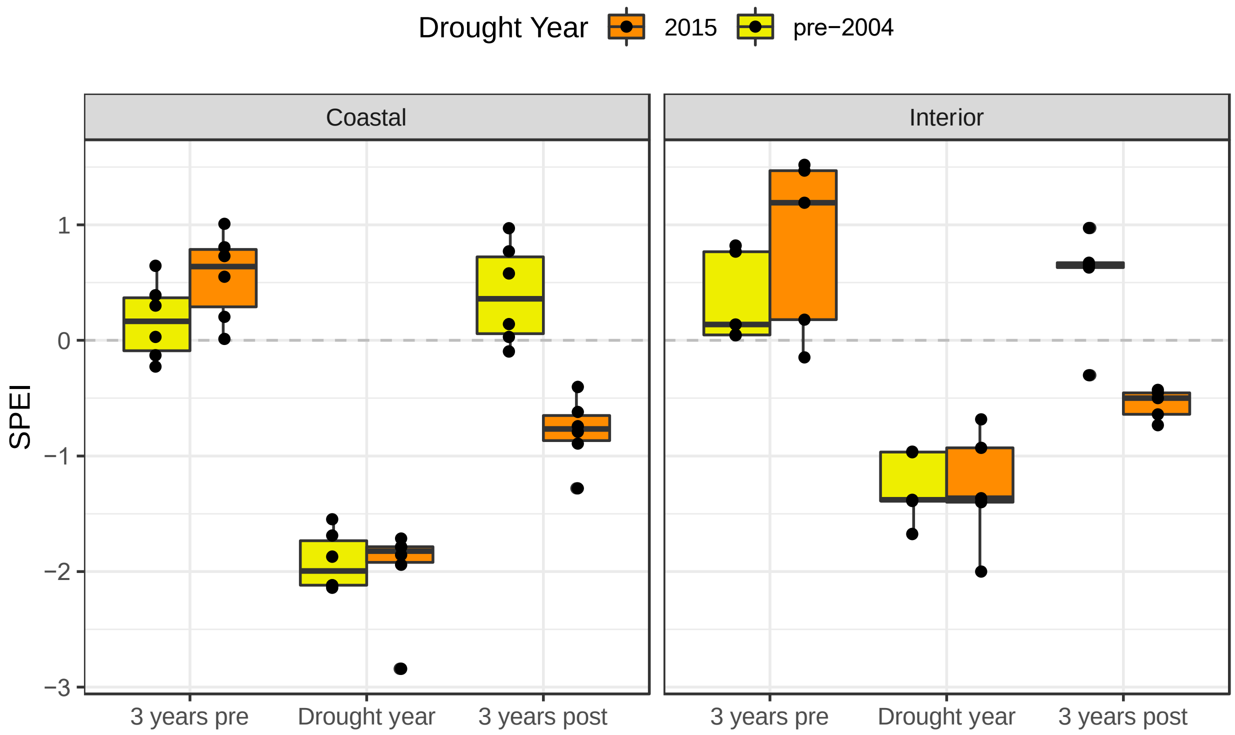

**Fig. S9:** May to June Standardized Precipitation Evaporation Index (SPEI) in the drought years, the average of site SPEI in the three years pre-drought, and the average of site SPEI in the three years post-drought in coastal (C) and interior (I) populations of WRC trees, illustrating similar SPEI pre-drought and during the drought, but notably warmer, drier conditions following the 2015 drought compared to the pre-2004 drought years. In the boxplots, the thick horizontal line within the box is the median, and the lower and upper hinges represent the interquartile range (IQR; 25th–75th percentiles) of the distribution. The whiskers extend +/- 1.5 times the IQR to form the limit for outliers.

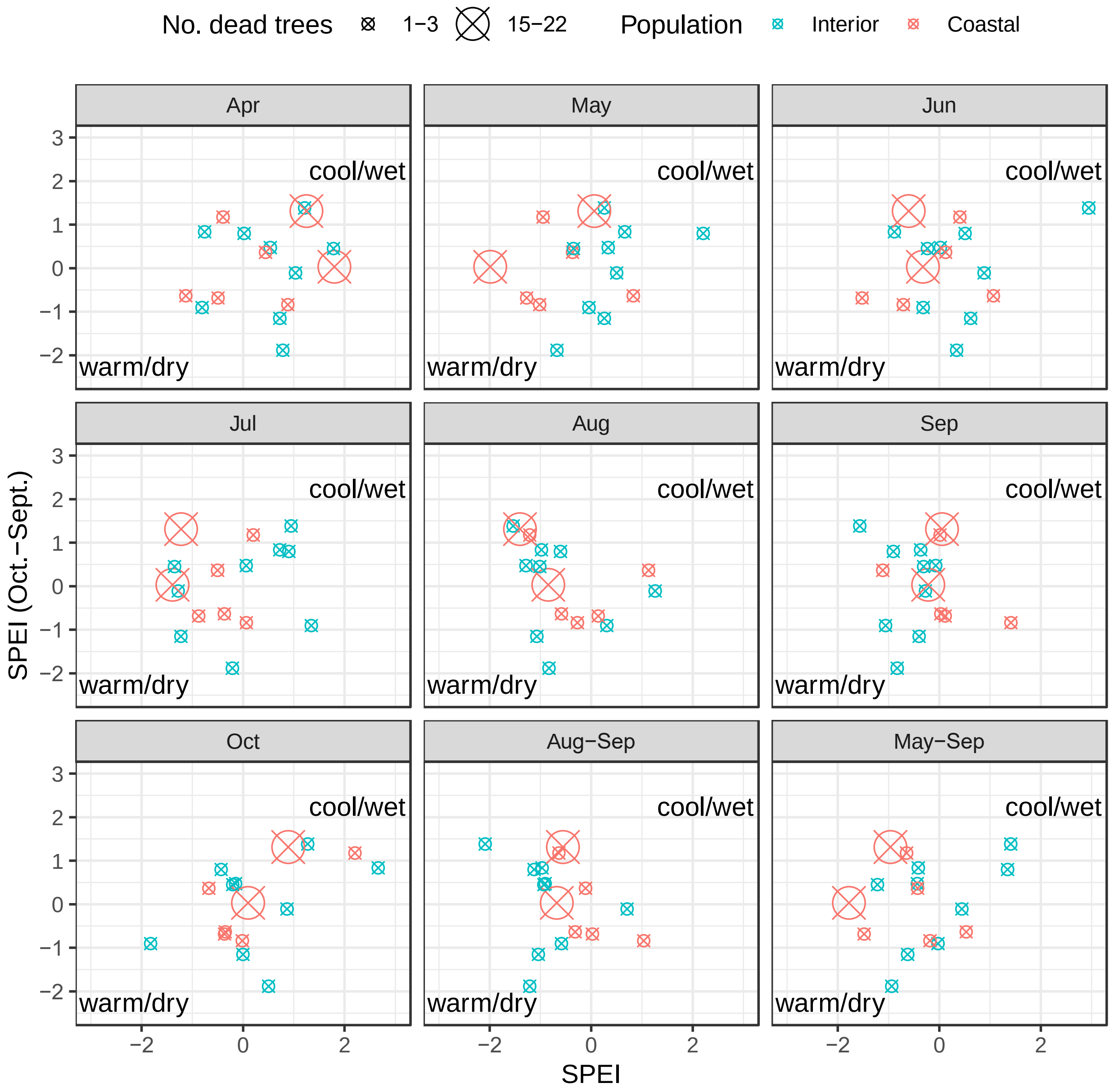


**Fig. S10:** Scatter plots of annual (y-axis) versus monthly, August-September, or May-September (x-axis) Standardized Precipitation Evaporation Index (SPEI) from 1970 to 2020 for interior and coastal populations. Circles with 'X' highlight years with mortality and size indicates number of dead trees sampled. SPEI was averaged across all sites within a population for each year. In coastal populations, mortality of 15-22 sampled trees (2 of 2 years and 80% of sampled mortality) occurred when May to September SPEI was ~ -1.0 or less (extremely warm/dry), and tree mortality of 1-3 sampled trees (4 of 5 years and 20% of sampled mortality) occurred when May to September SPEI was below average (warm/dry). Annual SPEI during these tree mortality years was average or positive (cool/wet). In interior populations, mortality occurred when August SPEI (7 of 9 years) and especially September SPEI (9 of 9 years) were average or negative (warm/dry), while annual SPEI during these years was both positive and negative. We selected May to September SPEI for comparison with annually-resolved death dates in coastal populations and August to September SPEI for comparison with annually-resolved death dates in interior populations.

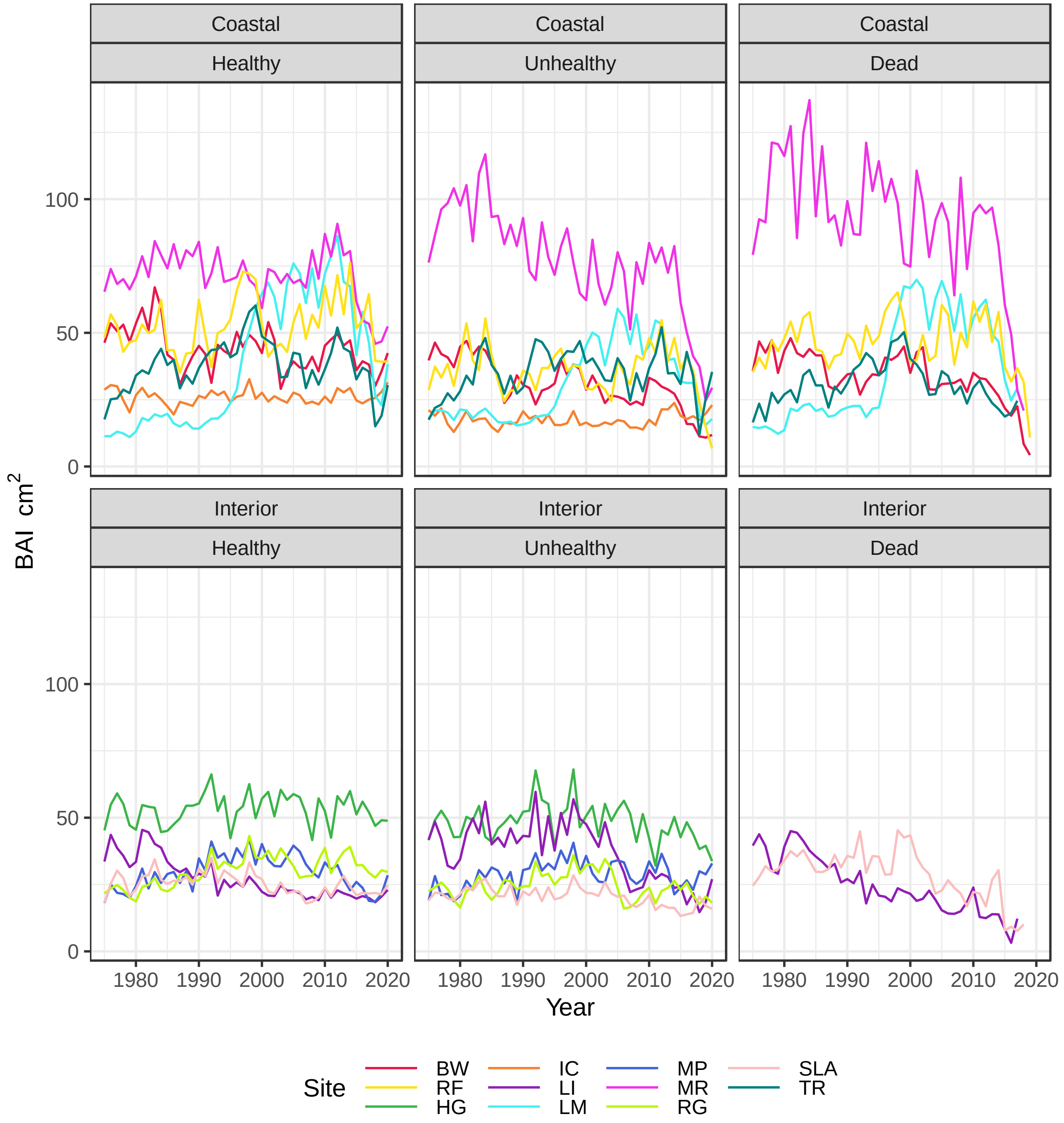


**Fig. S11:** Average basal area increment (BAI) by site for healthy (left), unhealthy (middle), and dead trees (right) in coastal (top) and interior (bottom) populations of western redcedar from 1975 to 2020.

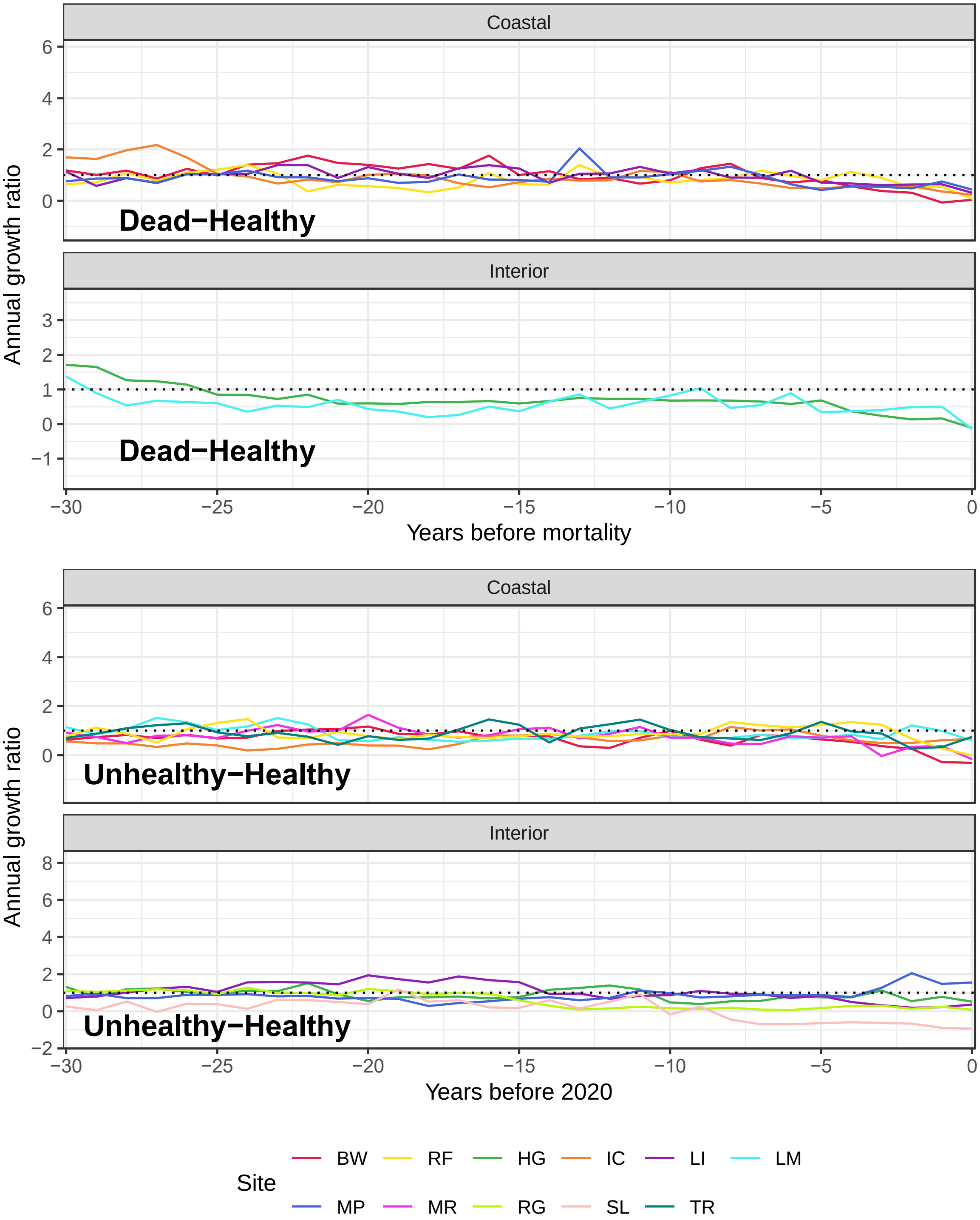


**Fig. S12:** Annual growth ratios for healthy vs. dead and healthy vs. unhealthy paired trees, calculated from the most recent 30 years of tree radial growth for coastal and interior populations. The black lines are site means, and the gray lines are individual tree pairs. An annual growth ratio significantly less than one indicates that healthy trees grew more than their paired, co-located dead or unhealthy trees in that year (or the opposite, if ratio >1). Site means of ratios for healthy vs. dead paired trees (top two panels) were visually similar (lines cross-cross and show similar trend) for interior and coastal populations. Site means of ratios for healthy vs. unhealthy paired trees (bottom two panels) were similar among coastal populations, but site means in interior populations exhibited different trajectories in the last ~10 years.

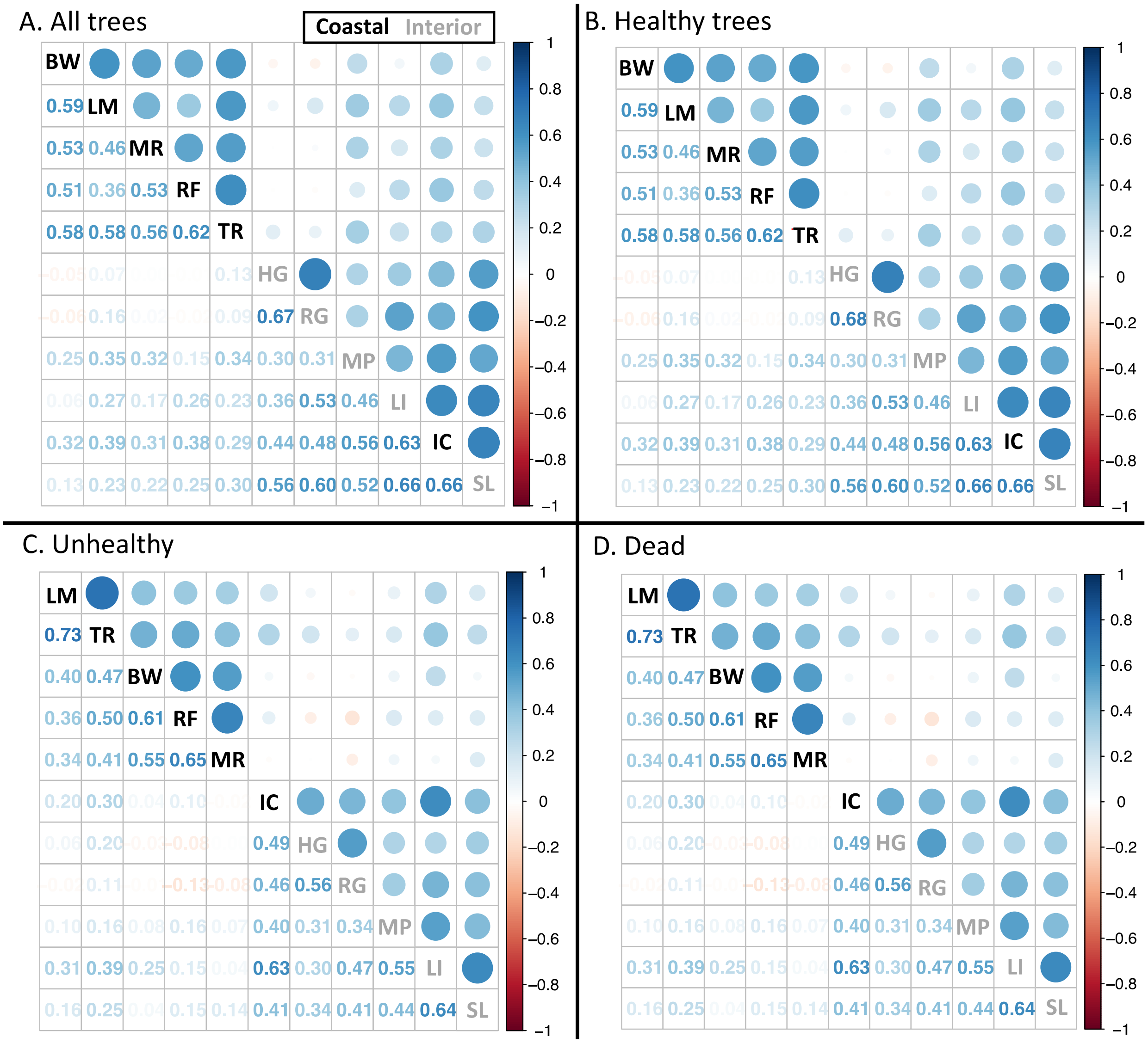

**Fig. S13:** Correlations table for site tree ring chronologies (1975-2020) for **A)** all trees, **B)** healthy trees, **C)** unhealthy trees, and **D)** dead trees. Sites are grouped with hierarchical clustering into coastal (black) and interior (gray) populations of western redcedar. 'IC', a coastal population site on the eastern side of the Cascade Mountains, was more strongly correlated with sites in interior populations. Circle size indicates strength of correlation coefficients, and the numerical values are Spearman's rho values.
